# Impaired Endosomal Recycling of Signaling Receptors Activates an Extracellular UPR

**DOI:** 10.64898/2026.03.12.711310

**Authors:** Avijit Mallick, Yunguang Du, Cole Haynes

## Abstract

Mitochondrial dysfunction and extracellular protein aggregation occur in neurodegenerative diseases such as Alzheimer’s disease (AD). However, it remains unclear if these processes are functionally linked. Here, we identify a signaling pathway that is activated upon accumulation of aggregation-prone proteins in the extracellular space. We find that the transcription factor ATFS-1, which regulates the mitochondrial unfolded protein response, also regulates transcripts required for endosomal recycling, multiple plasma membrane-localized signaling receptors, and secreted proteins that bind aggregation-prone proteins in the extracellular space, including transthyretin and Aβ, and promote their degradation. Interestingly, Aβ(1-42) aggregation induces *atfs-1*-dependent transcription by promoting degradation of the bZIP protein ZIP-3, which antagonizes ATFS-1. ZIP-3 accumulates in the cytosol when it is phosphorylated by kinases that function downstream of plasma membrane-localized signaling receptors, including the WNT and glutamate receptors. Upon ligand binding, the signaling receptors stimulate the cognate kinase, many of which we found phosphorylate ZIP-3, impeding ZIP-3 degradation, allowing it to antagonize *atfs-1*-dependent transcription. However, accumulation of aggregation-prone proteins such as Aβ(1–42) causes endosomal swelling, which impairs endosomal recycling, instead diverting signaling receptors to lysosomes for degradation. In turn, the depletion of signaling receptors reduces the level of ZIP-3 phosphorylation, resulting in ZIP-3 degradation and activation of *atfs-1*-dependent transcription, which promotes extracellular proteostasis. Our findings uncover an unexpected coupling between endocytic quality control and mitochondrial signaling, revealing a circuit that preserves extracellular proteostasis and promotes organismal resilience.

## INTRODUCTION

Canonical proteostasis pathways, including the endoplasmic reticulum unfolded protein response (UPR^ER^)^1^, heat shock response (HSR)^2^, and mitochondrial unfolded protein response (UPR^mt^)^3^, rely on organelle-specific sensors and transcription factors to restore local protein homeostasis; however, whether and how these systems detect and respond to proteostasis failure in the extracellular space remains unclear. Disruption of extracellular proteostasis is a defining feature of neurodegenerative diseases such as Alzheimer’s disease (AD), which is characterized by the accumulation of extracellular amyloid-β (Aβ) aggregates and widespread mitochondrial dysfunction^4–6^. Analyses of AD patient samples and disease models have revealed extensive metabolic reprogramming^7,8^, including increased glycolysis and activation of the UPR^mt^. Despite these observations, the mechanistic links between extracellular proteostasis perturbations and mitochondrial stress signaling remain poorly understood.

The UPR^mt^ is a mitochondria-to-nucleus signaling pathway that promotes mitochondrial recovery by activating transcription of numerous genes that promote mitochondrial biogenesis in response to mitochondrial stress^3^. In *Caenorhabditis elegans*, the UPR^mt^ is mediated by the bZIP transcription factor ATFS-1, which under basal conditions is mostly imported into mitochondria and degraded^9,10^. However, during mitochondrial stress, ATFS-1 fails to import into mitochondria and translocate to the nucleus, where it induces transcription of genes that promote mitochondrial proteostasis and biogenesis^9–11^. Importantly, ATFS-1 activity is also negatively regulated by the bZIP protein ZIP-3, which forms a heterodimer with cytosolic ATFS-1^12,13^. Inhibition of ZIP-3 expression is sufficient to activate ATFS-1-dependent transcription, independent of mitochondrial dysfunction.

Endosomal dysfunction is an early and prominent pathological feature of AD^14,15^. In patient samples and disease models, endosomal swelling disrupts the recycling of endocytosed cargo, such as membrane-localized signaling receptors, from endosomes back to the plasma membrane. Endosomal recycling is required to maintain plasma membrane composition, sustain receptor-mediated signaling, and preserve extracellular homeostasis^14,16–18^. In AD, accumulation of Aβ(1-42) within endosomes promotes endosomal swelling, impairing cargo recycling and diverting signaling receptors to lysosomes for degradation, thereby attenuating cell-surface signaling^14,15,19,20^.

The retromer complex, consisting of VPS26, VPS29, and VPS35, mediates endosomal recycling^16^, and perturbations in this pathway are causally implicated in AD^15,21^. Mutations in VPS35 are associated with familial forms of AD^15,21^ and Parkinson’s disease^22^ and have also been shown to cause mitochondrial dysfunction^23^, yet how defects in endosomal trafficking intersect with mitochondrial stress signaling remains unclear.

Here, we identify ATFS-1 as a regulator of extracellular proteostasis through transcriptional control of endosomal recycling and sorting components, as well as secreted extracellular chaperones and proteases that limit extracellular protein aggregation. We define a mechanism by which extracellular protein aggregation promotes ZIP-3 degradation at recycling endosomes, leading to ATFS-1 activation and induction of a transcriptional program that restricts extracellular aggregate accumulation and preserves plasma membrane receptor expression. ZIP-3 degradation is inhibited by phosphorylation mediated by kinases acting downstream of cell-surface receptors, including glutamate receptors. In contrast, disruption of endosomal recycling or receptor-mediated signaling promotes ZIP-3 degradation, thereby activating ATFS-1. Together, these findings define a stress-responsive signaling network in which endosomal trafficking, receptor signaling, and ATFS-1-dependent transcription maintain or restore extracellular proteostasis.

## RESULTS

### ATFS-1-dependent transcription impairs the accumulation of extracellular protein aggregates

We previously demonstrated that ATFS-1 promotes transcription of genes required for mitochondrial biogenesis, structure, and function. Interestingly, many genes encoding proteins secreted to the extracellular space are reduced in *atfs-1(null)* worms, suggesting that ATFS-1 also regulates transcription of genes that affect cellular activities beyond mitochondria (Supplementary Figure 1A-B, Supplementary Table 1). A previous study identified 57 genes that impair extracellular protein aggregation in *C. elegans*^24^. Interestingly, *atfs-1* is required to express many of these 57 extracellular protein aggregation regulators that promote extracellular proteostasis^24^ (Supplementary Figure 1A and 1D).

**Figure 1.**
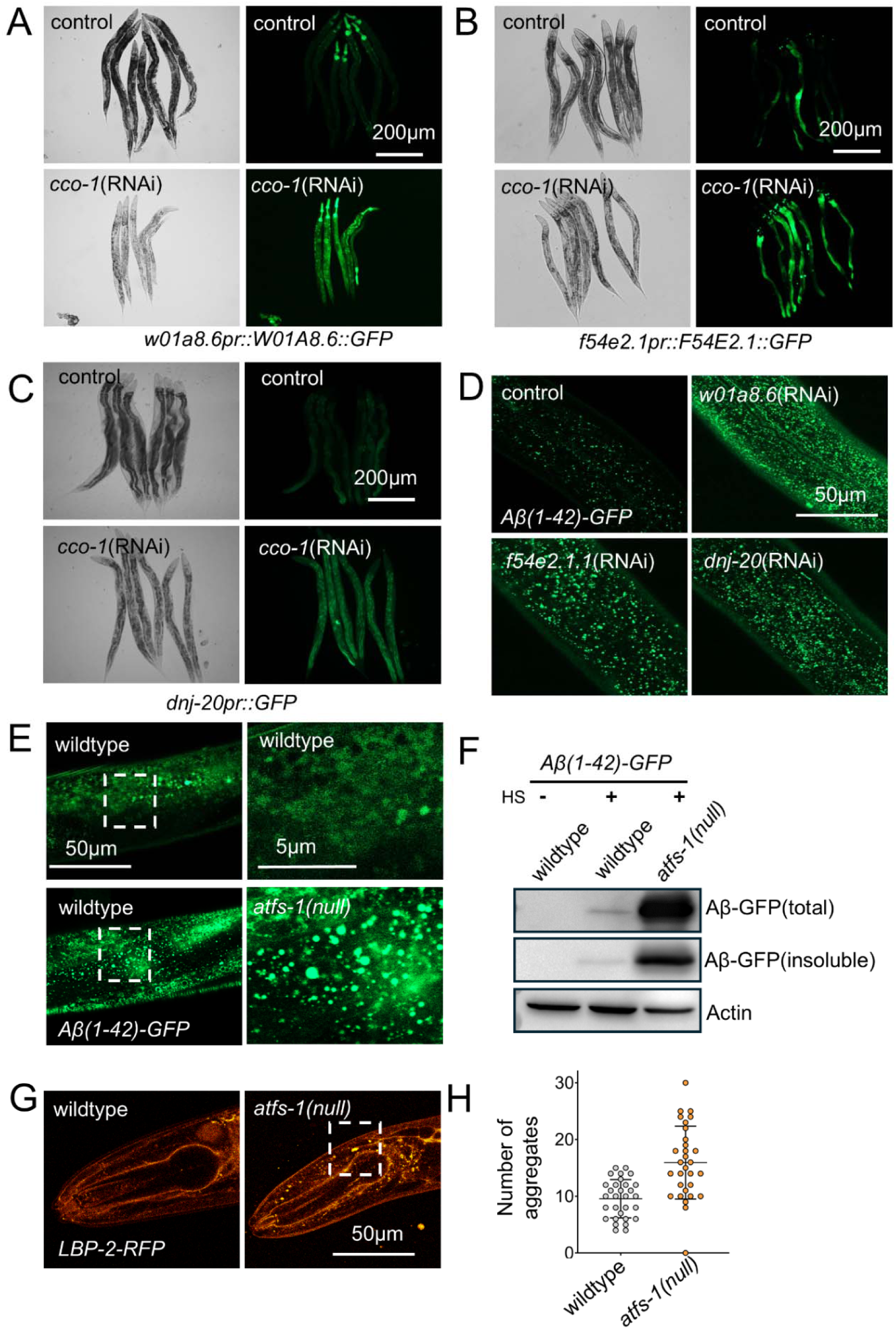
Activation of ATFS-1 promotes the expression of extracellular genes that limit aggregation. **A-C.** Representative images of worms expressing *W01A8.6_pr_::W01A8.6-GFP*, *F54e2.1_pr_::F54E2.1-GFP*, and *dnj-20_pr_::GFP* following control or *cco-1*(RNAi). Scale bar, 200μm. N =3, biologically independent replicates. **D.** Representative images of worms expressing *hsp-16.2_pr_::ssA*β*(1-42)-GFP* labelled protein aggregation puncta grown at 37°C for 1 hr during Day 1 adult following control, *W01A8.6*(RNAi), *F54E2.1*(RNAi), and *dnj-20*(RNAi). Scale bar, 50μm. N =3, biologically independent replicates. **E.** Representative images of worms expressing *hsp-16.2_pr_::ssA*β*(1-42)-GFP* labelled protein aggregation puncta grown at 37°C for 1 hr during Day 1 adult of wildtype and *atfs-1(null)* worms. Scale bar, 50μm. N =3, biologically independent replicates. **F.** Representative image of a Western blot analysis of total and insoluble fraction of *hsp-16.2_pr_::ssA*β*(1-42)-GFP* in wildtype and *atfs-1(null)* worms. **G.** Representative images of LBP-2-RFP labelled protein aggregation puncta of wildtype and *atfs-1(null)* Day 1 adults. Scale bar, 50μm. N =3, biologically independent replicates. **H.** Dot plots showing the number of LBP-2-RFP aggregation puncta, in wildtype and atfs-1(null) worms in **G**. N = 3, biologically independent samples. \*\*\**p* < 0.001 (two-tailed Student’s t-test).

To gain insight into the role of ATFS-1 in regulating these genes that promote extracellular proteostasis, we generated GFP-tagged reporter strains expressing *W01A8.6* (an extracellular metalloproteinase), *F54E2.1* (a membrane glycoprotein), and *dnj-20* (homolog of mammalian extracellular chaperone DNAJB11)^25^ (Figure 1A). RNAi inhibition of these genes resulted in elevated extracellular protein aggregates (Figure 1A-B). As ATFS-1 is required for the expression of *W01A8.6*, *F54E2.1,* and *dnj-20*, we hypothesized that the expression of these genes would respond to OXPHOS perturbation, which increases ATFS-1-dependent transcription^3,11,26^. As expected, *W01A8.6_pr_::GFP*, *F54E2.1_pr_::GFP,* and *dnj-20_pr_::GFP* transgenic strains exhibited increased GFP fluorescence when grown on RNAi targeting *cco-1* (complex IV gene) (Figure 1A-C).

To determine if *atfs-1* promotes extracellular proteostasis, we generated *atfs-1(null)* worms expressing either LBP-2::RFP or Aβ(1-42)-GFP. We find that *atfs-1(null)* worms have increased aggregation, indicating that ATFS-1 is required to impair extracellular protein aggregation (Figure 1E-H). In contrast, *cco-1* inhibition, which activates ATFS-1-dependent transcription, reduced the accumulation of LBP-2::RFP and Aβ(1-42)-GFP aggregates (Figure 2D and supplementary Figure 2E). Together, these findings indicate that *atfs-1* is required to impair extracellular protein aggregation.

**Figure 2.**
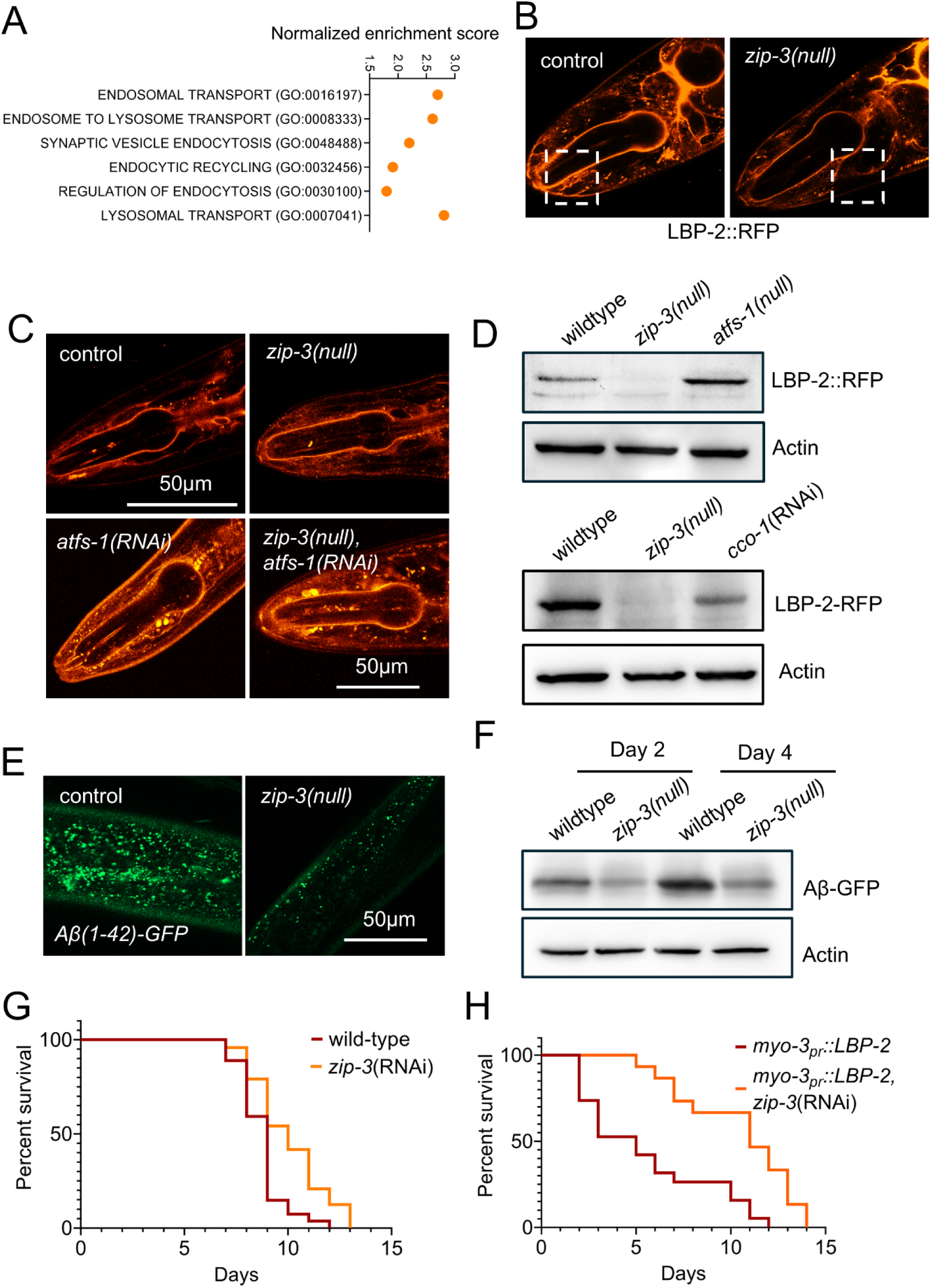
Inhibition of ZIP-3 reduces extracellular protein aggregation. **A.** Bubble plot showing the normalized enrichment scores of gene ontology classes for differentially expressed genes in *zip-3(null)* worms compared to wildtype. **B.** Representative images of LBP-2-RFP labelled protein aggregation puncta in wildtype and *zip-3(null)* Day 1 adults grown at 25°C. Scale bar, 50μm. N =3, biologically independent replicates. **C.** Representative images of LBP-2-RFP labelled protein aggregation puncta in wildtype, *zip-3(null), atfs-1(RNAi),* and *zip-3(null), atfs-1(RNAi)* Day 1 adults grown at 25°C. Scale bar, 50μm. N =3, biologically independent replicates. **D.** Representative image of a Western blot analysis of the insoluble fraction of *LBP-2-RFP* in wildtype, *zip-3(null)*, *atfs-1(null),* and *cco-1*(RNAi) worms. N =3, biologically independent replicates. **E.** Representative images of worms expressing *hsp-16.2_pr_::ssA*β*(1-42)-GFP* labelled protein aggregation puncta grown at 37°C for 1 hr during Day 1 adult of wildtype and *zip-3(null)* worms. Scale bar, 50μm. N =3, biologically independent replicates. **F.** Representative image of a Western blot analysis of the insoluble fraction of *hsp-16.2_pr_::ssA*β*(1-42)-GFP* in wildtype and *zip-3(null)* Day-2 and Day-4 adult worms. N =3, biologically independent replicates. **G-H.** Lifespan graphs for wildtype and *zip-3*(RNAi) worms grown at 25°C (**G**) and for *myo-3pr::LBP-2-GFP* worms grown at 20°C following control *and zip-3*(RNAi) (**H**). N =3, biologically independent replicates. \**p* < 0.01 (**G, H**) (log-rank test).

### Inhibition of ZIP-3 induces ECR transcription via ATFS-1 activation

We previously identified the bZIP protein ZIP-3 as a negative regulator of ATFS-1-dependent transcription^12^. Transcriptomic analyses confirmed that ZIP-3 represses transcription of ATFS-1 target genes while also regulating a broader transcriptional network independent of ATFS-1 and the mitochondrial unfolded protein response (UPR^mt^)^11,12^. Consistent with a functional interaction, prior studies demonstrated physical binding between ZIP-3 and ATFS-1^13^. Notably, ZIP-3–regulated genes were enriched for pathways involved in endosomal transport, endosome-to-lysosome trafficking, endocytic recycling, regulation of endocytosis, and lysosomal transport (Figure 2A).

To determine the impact of ZIP-3 on extracellular proteostasis, we examined aggregation of LBP-2::RFP and Aβ(1–42)::GFP in *zip-3(null)* worms. Interestingly, ZIP-3 deletion worms have reduced aggregates than wild-type animals (Figure 2E-F), even under elevated temperature stress at 25°C (Figure 2B). In contrast, inhibition of *atfs-1* increased aggregation in *zip-3*–deletion animals, suggesting that *atfs-1* is required to promote extracellular proteostasis in *zip-3(null)* worms (Figure 2C-D). Consistent with improved extracellular protein aggregation, *zip-3*(RNAi) also extended lifespan at 25°C (Figure 2G).

To further assess ZIP-3 function in extracellular proteostasis, we generated a transgenic strain expressing LBP-2::GFP levels under the muscle myosin (*myo-3*) promoter. We confirmed that LBP-2 is secreted into the pseudocoelomic space, internalized by coelomocytes, and forms aggregates by day 1 of adulthood, coinciding with reduced lifespan (Supplementary Figure 2A-B). The level of aggregation is markedly enhanced compared to the *lbp-2_pr_::lbp-2-RFP* (LBP-2-RFP) strain used before. Strikingly, *zip-3*(RNAi) reduced LBP-2::GFP aggregation and extended lifespan (Figure 2H and Supplementary Figure 2A). Supporting these findings, RNA-seq analysis revealed that *zip-3(null)* mutants increased the expression of genes involved in endocytosis and endosomal recycling, including 52 genes encoding secreted extracellular proteins such as collagens, oxidoreductases, proteases, and protease inhibitors (Supplementary Table 2). Among these are seven genes previously shown to limit extracellular aggregation^24,27^, which are reduced in *atfs-1(null)* mutants but elevated in *zip-3* deletion mutants (Supplementary Figure 1E-F). Together, these results demonstrate that ZIP-3 deletion enhances ATFS-1–dependent extracellular proteostasis, linking transcriptional activation of genes required to limit extracellular protein aggregation, promote endosomal trafficking, and improve organismal survival.

### ATFS-1 activation prevents the accumulation of extracellular protein aggregates and limits neurodegeneration

To assess the physiological relevance of ZIP-3 in regulating extracellular proteostasis, we used transgenic *C. elegans* models that secrete either human transthyretin (TTR) or Aβ(1–42) into the extracellular space. To model TTR-associated proteotoxicity, we used the aggregation-prone TTR(V30M) variant under the *unc-54* promoter, while Aβ(1–42) is expressed in either neuronal or muscle tissues. In both models, the secreted proteins accumulated in the extracellular space, providing robust systems to examine mechanisms governing extracellular protein quality control^28,29^.

We evaluated functional outcomes, including lifespan, body bending, and thrashing behavior. RNAi-mediated inhibition of *zip-3* significantly rescued motor dysfunction and extended lifespan in both Aβ(1–42)- and TTR(V30M)-secreting animals (Figure 3A-D and supplementary Figure 2C-D). Given that *zip-3* inhibition and ATFS-1 activation are each associated with improved healthspan, we next examined whether extracellular aggregation could activate ATFS-1–dependent transcription. We found that the neuronal and muscle Aβ(1–42)-expressing strains and TTR(V30M) worms exhibited increased *hsp-6p::GFP* expression, indicative of ATFS-1–dependent transcriptional activation (Figure 3E). Consistent with this, neuronal and muscle-secreted Aβ worms have reduced ZIP-3 levels (Figure 3G-H). Moreover, multiple genes that limit extracellular protein aggregation and have a reduced expression in *atfs-1(null)* animals are found to be induced in both Aβ(1–42) and TTR(V30M) transgenic strains (Supplementary Figure 1J-K).

**Figure 3.**
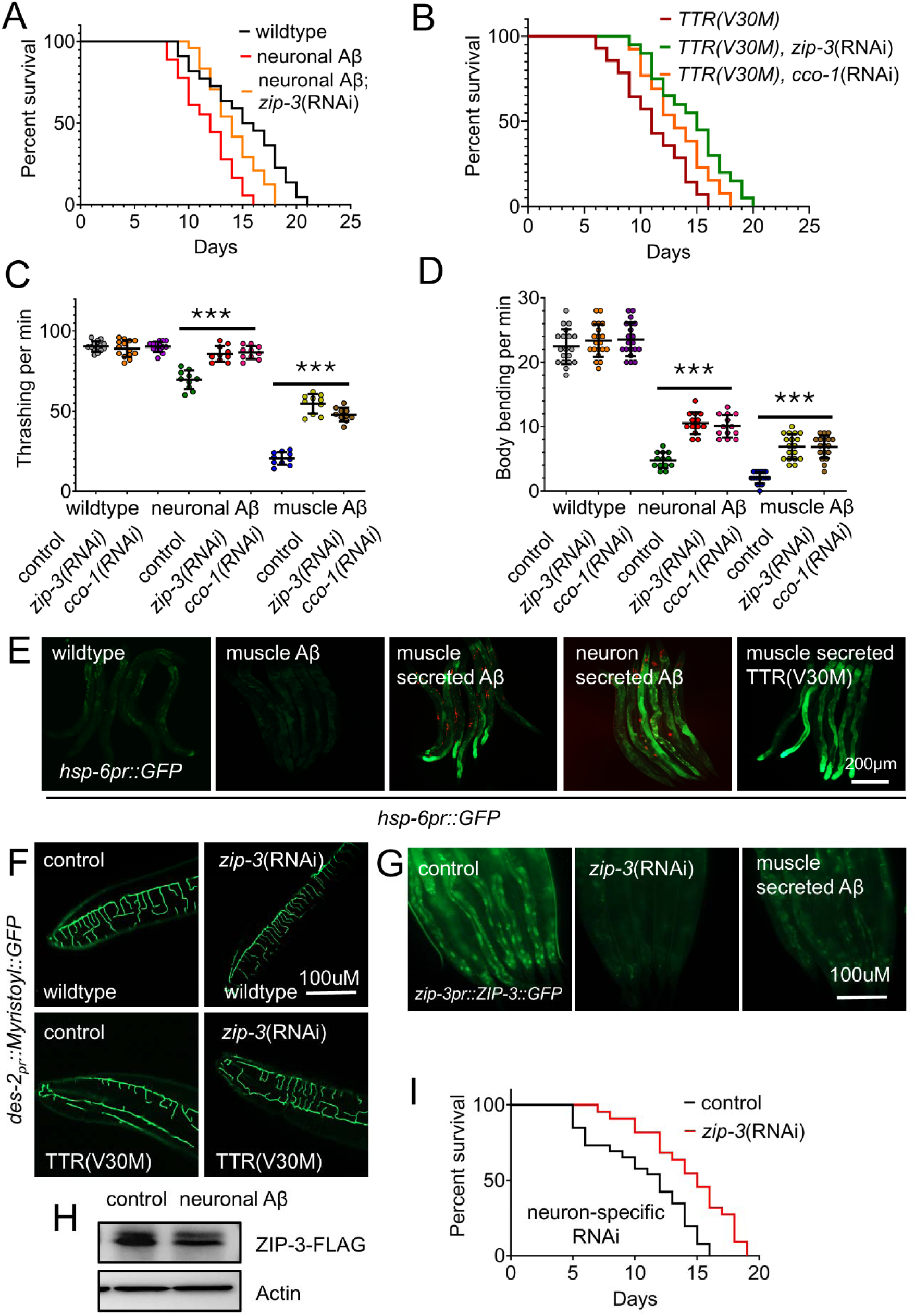
Inhibiting *zip-3* and the ETC complex component *cco-1* ameliorates phenotypes in extracellular protein aggregation models. **A-B.** Lifespan graphs for wildtype, neuronal Aβ(1-42) (**A**), and TTR(V30M) (**B**) worms grown following control *and zip-3*(RNAi) (**H**). N =3, biologically independent replicates. \**p* < 0.01 (**A, B**) (log-rank test). **C-D** Dot plots showing the rate of thrashing and body bending of neuronal- and muscle secreted-Aβ(1-42) worms grown on control, *zip-3*(RNAi), and *cco-1*(RNAi) plates. Data represents the mean of three independent biological replicates with error bars showing the standard error of the mean. N = 3, biologically independent samples. \*\*\**p* < 0.001 (one-way ANOVA). **E.** Representative image of *hsp-6_pr_::GFP* expression in wildtype, muscle Aβ (not secreted), muscle secreted Aβ(1-42), neuron secreted Aβ(1-42), and muscle secreted TTR(V30M). Scale bar, 200μm. N =3, biologically independent replicates. **F.** Representative images of the dendritic morphology of FLP neurons marked with *des-2_pr_::Myristoyl::GFP* of wildtype and TTR(V30M) worms following control and *zip-3*(RNAi). Scale bar, 100μm. N =3, biologically independent replicates. **G.** Representative image of *zip-3_pr_::ZIP-3::GFP* expression in wildtype, *zip-3*(RNAi) and muscle secreted Aβ(1-42) worms. Scale bar, 100μm. N =3, biologically independent replicates. **H.** Representative image of a western blot analysis of ZIP-3-3x-FLAG in wildtype and neuron-secreted Aβ(1-42) worms. N =3, biologically independent replicates. **I.** Lifespan graph of neuron-specific RNAi strain (MAH677) grown on control and *zip-3*(RNAi). N =3, biologically independent replicates. \**p* < 0.01 (log-rank test).

To directly assess neurodegenerative phenotypes in the presence of extracellular aggregation, we utilised the FLP neuronal marker *des-2p::Myristoyl::GFP* in the TTR(V30M) strain, shown to have impaired nociception and defective dendritic morphology^28^. We find that inhibiting *zip-3* improved the dendritic morphology in these worms (Figure 3F). We also examined GABAergic and dopaminergic neurons using *unc-47_pr_::GFP* and *dat-1_pr_::GFP* reporters^30,31^. Both *zip-3* overexpression and *atfs-1* deletion enhanced neuronal puncta formation by day 1 of adulthood, consistent with accelerated neurodegeneration (Supplementary Figure 2F-G). Consistent with this protective role of ATFS-1 and the expression of ZIP-3 and ATFS-1 in the head (Supplementary Figures 1C and 1L), inhibition of *zip-3* specifically in the neurons extended the lifespan of worms (Figure 3I). Together, these results show that extracellular proteotoxic stress triggers an ATFS-1–dependent transcriptional program that reduces extracellular aggregation, a process otherwise inhibited by ZIP-3.

### Endocytic recycling is required to limit extracellular aggregates and maintain ZIP-3 levels

Endosomal recycling is essential for maintaining plasma membrane integrity and extracellular protein quality control. Also, impaired endosomal recycling is a common pathology in AD progression. To determine whether Aβ(1–42) expressing worms impact the endosomal recycling, we utilised the late endosomal marker RAB-7::GFP. Expression of Aβ(1–42) in *C. elegans* resulted in a significant enlargement of RAB-7::GFP-labeled late endosomes (Figure 4A). This phenotype aligns with the endocytic trafficking impairments typically induced by extracellular proteotoxic stress^15^. Knockdown of *vps-35*, a core component of the retromer complex, further exacerbated endosomal swelling, indicating that impaired retromer-dependent recycling intensifies endosomal dysfunction (Figure 4A). Interestingly, *atfs-1* mutants exhibited disrupted endosomal morphology both basally and following *vps-35* depletion (Figure 4B), suggesting that ATFS-1 is required to preserve endosomal architecture. Notably, RNAi-mediated inhibition of *zip-3* or *cco-1* reduced the endosomal enlargement in Aβ(1–42)–expressing worms (Figure 4C), consistent with a protective role of ATFS-1 activation in maintaining endosomal homeostasis under extracellular proteotoxic stress.

**Figure 4.**
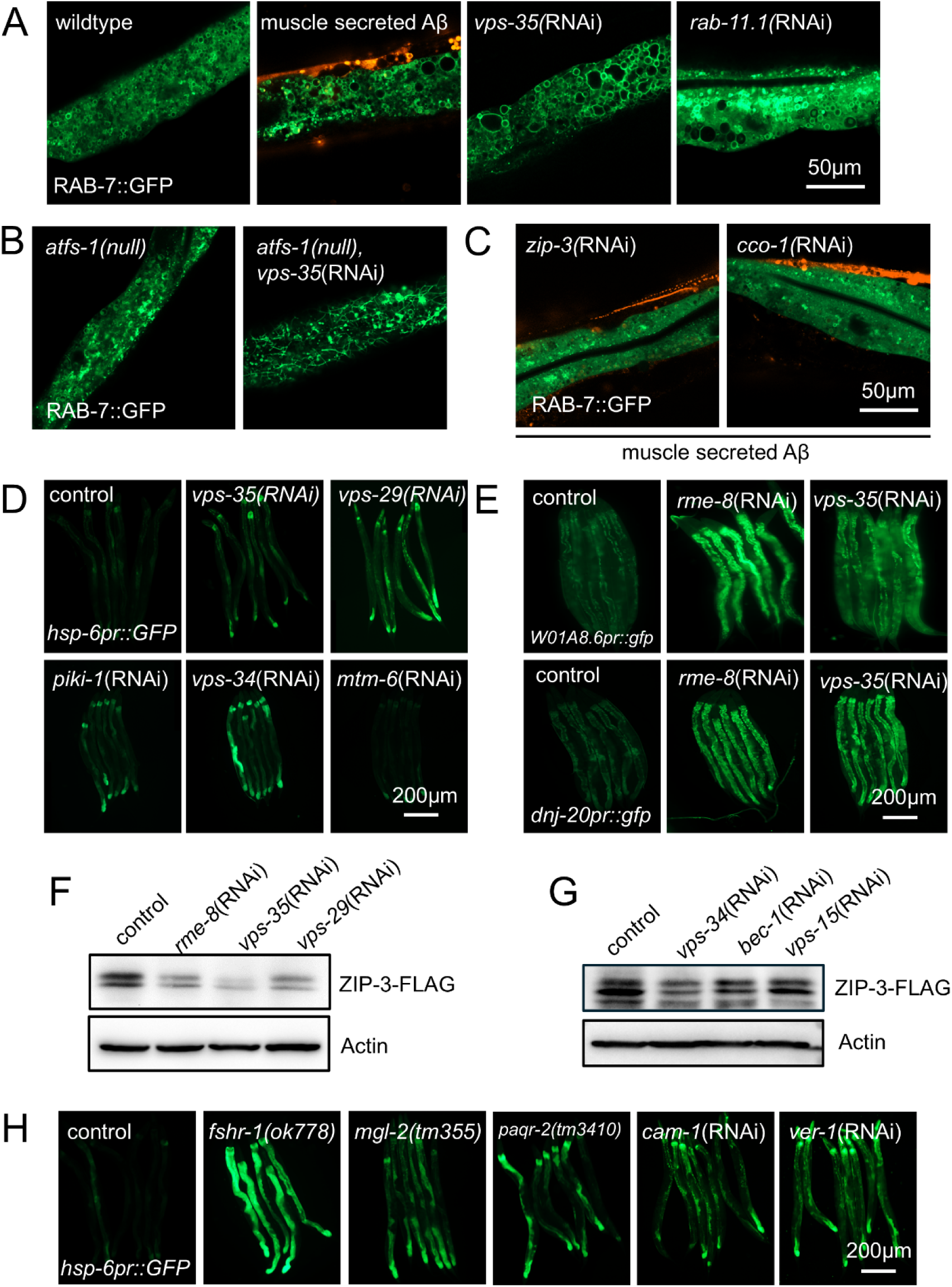
Inhibition of endosomal recycling leads to reduced ZIP-3 level and expression of extracellular genes dependent on ATFS-1 activation. **A.** Representative images of late endosomal marker RAB-7::GFP in wildtype, muscle secreted Aβ(1-42), *vps-35*(RNAi), and *rab-11.1*(RNAi). **B.** Representative images of late endosomal marker RAB-7::GFP in *atfs-1*(null) worms grown on control and *vps-35*(RNAi) HT115 bacteria. **C.** Representative images of late endosomal marker RAB-7::GFP in muscle secreted Aβ(1-42) worms grown on *zip-3*(RNAi) and *cco-1*(RNAi) HT115 bacteria. Enlarged endosomes seen in muscle secreted Aβ(1-42) worms (**A**) are rescued following *zip-3*(RNAi) and *cco-1*(RNAi) (**C**). Scale bar, 50μm. N =3, biologically independent replicates. **D.** Representative image of *hsp-6_pr_::GFP* expression following control RNAi, *vps-35*(RNAi), *vps-29*(RNAi), *piki-1*(RNAi), *vps-34*(RNAi) and *mtm-6*(RNAi). Scale bar, 200μm. N =3, biologically independent replicates. **E.** Representative image of extracellular protease *w01a8.6_pr_::GFP* and extracellular chaperone *dnj-20_pr_::GFP* expression following control RNAi, *vps-35*(RNAi), and *rme-8*(RNAi). Scale bar, 200μm. N =3, biologically independent replicates. **F-G.** Representative image of a western blot analysis of ZIP-3-3x-FLAG following inhibition of endosomal retromer complex components (*rme-8, vps-35*, and *vps-29*) and class III PI3K components (*vps-34, bec-1,* and *vps-15*). N =3, biologically independent replicates. **H.** Representative image of *hsp-6_pr_::GFP* expression in wildtype, *fshr-1(ok778), mgl-2(tm355), paqr-2(tm3410)*, *cam-1*(RNAi) and *ver-1*(RNAi). Scale bar, 200μm. N =3, biologically independent replicates.

The retromer complex, composed of VPS-26, VPS-29, and VPS-35, coordinates endosomal recycling to the plasma membrane^16,17^ and retrograde trafficking to the trans-Golgi network via RME-8 and HSP-1^17^. To test whether disruption of these pathways activates ATFS-1, we individually inhibited the expression of retromer components (*vps-26, vps-29,* and *vps-35*), retrograde regulators (*rme-8* and *hsp-1*), and additional endosomal recycling-associated factors (*rme-1, sdpn-1, alx-1, rab-10, rab-11.1,* and *rab-35*)^32^. Each perturbation robustly induced the ATFS-1 reporter *hsp-6_pr_::GFP* (Figure 4D). Consistent with ATFS-1–dependent extracellular proteostasis, inhibition of *vps-26, vps-35,* or *rme-8* also induced expression of extracellular genes, including the protease *W01A8.6_pr_::GFP* and the chaperone *dnj-20_pr_::GFP* (Figure 4E and supplementary Figure 1G-I). Importantly, mitochondrial membrane potential remained intact following *rme-8* knockdown, indicating that ATFS-1 activation in this context occurs independently of overt mitochondrial dysfunction (Supplementary Figure 3B).

To understand whether ATFS-1-dependent transcription is specific to perturbation in the retromer complex or in general related to the health of endosomal recycling, we perturbed the phosphatidylinositol-3-phosphate [PI(3)P] synthesis pathway. PI(3)P is a key lipid signal that defines early endosomes and drives endosomal recycling and is generated primarily by class III PI3-kinase VPS-34, in complex with VPS-15 and BEC-1^32,33^. Consistent with a role for PI(3)P in endosomal recycling^34^, RNAi-mediated knockdown of *vps-34*, as well as *piki-1* (phosphoinositide-3-kinase), robustly induced *hsp-6p::GFP* expression (Figure 4D). In contrast, inhibition of the PI(3)P phosphatase *mtm-6*, which increases PI(3)P levels, did not induce *hsp-6_pr_::GFP* expression (Figure 4D). We next examined whether disruption of endosomal trafficking activates ATFS-1 through ZIP-3 destabilization. ZIP-3::GFP abundance was significantly reduced following knockdown of *rme-8, vps-26, vps-29,* or *vps-35*, a result confirmed in animals expressing endogenously tagged ZIP-3–FLAG (Figure 4F and supplementary Figure 3A). Similarly, depletion of *vps-34* caused enlargement of early and late endosomes and decreased ZIP-3 levels, indicating that PI(3)P-dependent trafficking and retromer function are required to maintain ZIP-3 stability under homeostatic conditions (Figure 4G). Together, these results demonstrate that disruption of endosomal recycling, through impaired retromer activity or PI(3)P synthesis, destabilizes ZIP-3, relieves repression of ATFS-1, and activates the unfolded protein response in the extracellular compartment to restore extracellular proteostasis.

### Endosomal recycling of plasma membrane-localized receptors promotes ZIP-3 stabilization via phosphorylation

Inhibition of endosomal recycling reduced ZIP-3 levels and activated ATFS-1–dependent transcription, suggesting that diminished cell surface receptor signaling initiates this response. To identify receptors regulating ZIP-3, we performed an RNAi screen targeting plasma membrane receptors and monitored ATFS-1 activity using the *hsp-6_pr_::GFP* reporter. Knockdown of multiple G protein–coupled receptors (GPCRs), including *fshr-1, mgl-2, npr-28,* and *paqr-2*, as well as non-GPCR receptors such as *cam-1* (Ror), *lin-12* (Notch), *lin-17* (Wnt Frizzled)*, vab-1* (Ephrin), and *ver-1* (EGFR), induced *hsp-6* expression (Figure 4H). WWP-1 recognizes a specific consensus binding motif, PPxY, on its substrates^35^, facilitating efficient internalization and endocytic recycling of the receptors^36^ (Supplementary Figure 3G). Notably, all non-GPCR receptors that induced *hsp-6_pr_::GFP* when inhibited using RNAi contained PPxY motifs. To determine whether the absence of interaction between the receptors and WWP-1 activates ATFS-1-dependent transcription, we mutated the specific PPxY sites to PPxA on the *cam-1* and *ver-1* receptors. Impressively, mutating the PPxY sites on *cam-1* and *ver-1* induces *hsp-6::GFP* (Supplementary Figure 3F), suggesting that the ZIP-3 turnover and activation of ATFS-1-dependent transcription initiates with the lack of WWP-1 interaction with the cell surface receptors. This activation of ATFS-1-dependent transcription is suppressed in ZIP-3(YA) mutants that lack the PPxY motif owing to the lack of ZIP-3 degradation by WWP-1 (Supplementary Figure 3H). Moreover, ZIP-3::GFP colocalized with the recycling endosome marker RAB-11.1::mRuby upon WWP-1 inhibition (Figure 5B), and WWP-1 depletion induced tubular ZIP-3::GFP structures (Figure 5A). This data suggests that the WWP-1-mediated ZIP-3 turnover takes place on the endosomal membrane. Similar to increased ZIP-3 accumulation, stalled tubular endosomal morphology is observed in *atfs-1(null)* worms following *vps-35* knockdown (Figure 4B), suggesting that ATFS-1 is required to sustain endosomal recycling.

**Figure 5.**
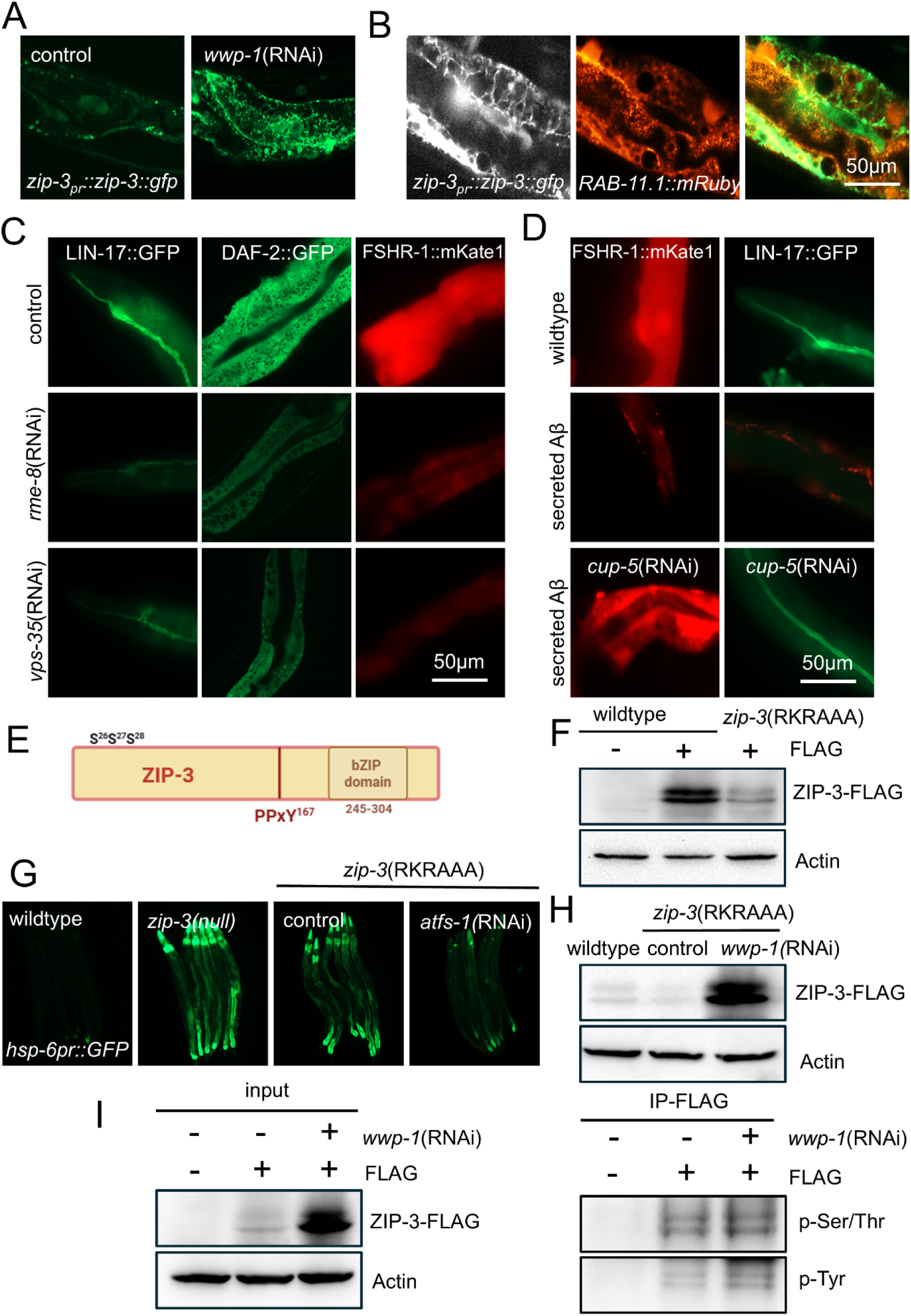
Kinases downstream of cell signaling receptors that undergo endosomal recycling regulate ZIP-3 via phosphorylation. **A.** Representative image of *zip-3_pr_::ZIP-3::GFP* expression following control and *wwp-1*(RNAi). Scale bar, 50μm. N =3, biologically independent replicates. **B.** Representative image showing overlay of *zip-3_pr_::ZIP-3::GFP* and endosomal recycling marker RAB-11.1::mRuby expression following *wwp-1*(RNAi). Scale bar, 50μm. N =3, biologically independent replicates. **C.** Representative images of cell signaling receptors LIN-17/Frizzled, DAF-2/Insulin-like receptor, and FSHR-1/GPCR expression following inhibition of *rme-8* and *vps-35*. **D.** Representative images of FSHR-1 and LIN-17 expression in wildtype and muscle secreted Aβ(1-42) worms following control and *cup-5*(RNAi). **C-D.** Scale bar, 50μm. N =3, biologically independent replicates. **E.** Schematic of ZIP-3 protein showing the conserved phosphorylation site RKRSSS (26-28aa), PPxY motif (167aa), and bZIP domain (245-304aa). **F.** Representative image of a western blot analysis of ZIP-3-3x-FLAG in wildtype worms lacking the FLAG tag and with the FLAG tag, and in ZIP-3(RKRAAA) mutant with the RKRSSS site mutated. N =3, biologically independent replicates. **G.** Representative image of a western blot analysis of ZIP-3-3x-FLAG in wildtype and ZIP-3(RKRAAA) mutant following control RNAi and *wwp-1* inhibition. N =3, biologically independent replicates. **I.** Representative image of a western blot analysis of ZIP-3-3x-FLAG after coimmunoprecipitation (Co-IP) in wildtype worms lacking the FLAG tag and with the FLAG tag following control RNAi and *wwp-1* inhibition. Phosphotyrosine and phosphoserine/threonine antibodies are used to blot for phosphorylation of tyrosine, serine, and threonine residues on ZIP-3. N =3, biologically independent replicates.

Because GPCRs lack PPxY motifs, we examined arrestin adaptor proteins that harbor PPxY motifs and mediate WWP-1-dependent endocytosis of GPCR^36^. RNAi depletion of *arrd-8, arrd-10, arrd-11,* and *arrd-22*, as well as disruption of G protein signaling via the Gα subunit *gsa-1*, induced *hsp-6* (Supplementary Figure 3D), implicating GPCR-mediated signaling in the regulation of ZIP-3. Consistent with impaired endosomal recycling leading to reduced receptor expression, inhibition of *vps-35* or *rme-8* reduced the abundance of fluorescently tagged FSHR-1, LIN-17, and DAF-2 receptor expression and concomitantly decreased ZIP-3 levels (Figure 5C and supplementary Figure 3A and 3E). We also observed a reduction in the receptor expression in the Aβ(1–42) expressing strain that showed endosomal swelling similar to *vps-35* inhibition (Figure 5D). During normal endosomal recycling, receptors are trafficked back to the cell surface membrane via retromer-dependent endosomal recycling or direct endosomal recycling. However, if the recycling pathway is perturbed, endosomal cargoes are directed to lysosomes for degradation. To test whether inhibiting lysosomal uptake can block the reduction in receptor expression, we inhibited *cup-5* in Aβ(1–42) expressing worms. CUP-5 is a mammalian orthologue of MCOLN1 expressed on the endo-lysosomal and lysosomal membranes and is required for lysosomal uptake and function^37^. We found that *cup-5* inhibition in the Aβ(1–42) worms reverted and increased FSHR-1 and LIN-17 expression (Figure 5D), suggesting that during perturbed endosomal swelling, receptors are degraded in the lysosomes, which activates ATFS-1-dependent transcription. Supporting our model of UPR^EC^, we found that inhibiting the receptor expression (*fshr-1*, *lin-17*, and *cam-1*) reduced protein aggregation in the Aβ(1–42) worms by activating ATFS-1 dependent transcription (Supplementary Figure 4A).

Because ZIP-3 degradation followed receptor loss, we determined whether ZIP-3 stability depends on phosphorylation by kinases downstream of cell surface receptor signaling. Our bioinformatics analysis shows that ZIP-3 contains multiple predicted phosphorylation sites (Figure 5E and supplementary Figure 4B). We used CRISPR-Cas9 to generate site-directed mutagenesis of a conserved phosphorylation cluster on ZIP-3 (RKRSSS→RKRAAA)^38^. This decreased ZIP-3 levels and induced *hsp-6_pr_::GFP* expression similar to the *zip-3* deletion mutant (Figure 5F-G). ZIP-3 abundance in this mutant is restored by inhibiting *wwp-1*, indicating that WWP-1 selectively targets the dephosphorylated ZIP-3 protein (Figure 5H). Co-immunoprecipitation of ZIP-3–FLAG followed by phospho-specific immunoblotting revealed phosphorylation on both serine and tyrosine residues on ZIP-3 (Figure 5I). This is also supported by an RNAi screen of families of kinases that are predicted to have conserved phosphorylation sites on ZIP-3, which identified *kin-1* (PKA), *pkc-2* (PKC), *akt-2* (AKT), and *kgb-1* (JNK) as kinases required to maintain ZIP-3 phosphorylation and stability (Supplementary Figure 4B-D).

WWP-1 activity is regulated by a conserved C2 domain that mediates cell membrane association and WWP-1 self-autoinhibition through interactions with the HECT domain^39,40^. To test whether disruption of this domain alters WWP-1 activity, we targeted the predicted C2 region of *wwp-1* using CRISPR–Cas9 (Supplementary Figure 5B). While complete deletion of this region was not viable, we isolated a mutant containing an insertion of two amino acids (N, E) within the C2 domain (*wwp-1(NE)*) (Supplementary Figure 5A-B). This mutant exhibited elevated *hsp-6* expression compared to *wwp-1(null)* animals, consistent with a partial gain-of-function phenotype, and showed extended lifespan, in contrast to the reduced viability of *wwp-1(null)* mutants (Supplementary Figure 5A-D). Together, these results demonstrate that surface receptor signaling and endosomal recycling maintain ZIP-3 stability through phosphorylation of ZIP-3 and C2 domain–regulated WWP-1 activity. In contrast, impaired endosomal trafficking or loss of cell signaling receptors promotes WWP-1-mediated ZIP-3 degradation, relieving inhibition of ATFS-1 by ZIP-3 and activating a transcriptional program that limits extracellular protein aggregation.

### Mitochondrial perturbations that extend lifespan also promote EC proteostasis

Previous studies have established that mild inhibition of mitochondrial electron transport chain (ETC) components, such as *nuo-6*, *isp-1*, and *cco-1*, extends lifespan via an ATFS-1-dependent transcriptional program^26,41–43^. However, the precise downstream mechanisms translating this stress response into longevity remain elusive. We hypothesized that ATFS-1 may promote organismal survival by promoting extracellular proteostasis. Through a targeted screen for ATFS-1-dependent chaperones, we identified *dnj-20*, *W01A8.6*, and *F54E2.1* as a critical effector. Knockdown of *dnj-20, W01A8.6,* and *F54E2.1* resulted in a marked accumulation of Aβ(1–42)-positive aggregates (Figure 1D). Furthermore, chromatin immunoprecipitation confirmed that ATFS-1 directly binds the *dnj-20* promoter, establishing *dnj-20* as a primary transcriptional target likely responsible for stabilizing the extracellular proteome under mitochondrial stress.

To determine the functional necessity of this extracellular proteostasis axis for the longevity of *isp-1(qm150)* and *clk-1(qm30)* mutants, we inhibited *dnj-20* and *W01A8.6*. Notably, the knockdown of *dnj-20*, or the deletion of *W01A8.6*, significantly suppressed the extended lifespan of these OXPHOS mutants (Supplementary Figure 6A-C). Together, these findings establish the ATFS-1–mediated extracellular unfolded protein response (UPR^EC^) as an essential regulator of proteostasis and longevity. Our work uncovers a cell-non-autonomous signaling mechanism that aligns stress responses with extracellular protein quality control to sustain organismal health and lifespan.

## Discussion

Although proteostasis has largely been examined through intracellular stress pathways, the maintenance of the extracellular (EC) proteome is equally important for tissue homeostasis and intercellular communication^44,45^. Here, we define an extracellular unfolded protein response (UPR^EC^) centered on ATFS-1 that is activated by inhibition or degradation of the bZIP transcription factor ZIP-3. Inhibition of ZIP-3 activates an ATFS-1–dependent transcriptional program that coordinates extracellular proteostasis through promoting expression of genes encoding extracellular chaperones, extracellular proteases, cell surface receptors, and endosomal recycling components. These findings position ATFS-1 as a central transcription factor linking extracellular protein quality control to endosomal trafficking and nuclear gene regulation.

A key advance of this study is the demonstration that perturbations in endosomal recycling, an early and conserved feature of Alzheimer’s disease (AD) pathology, directly activate UPR^EC^ through destabilization of ZIP-3. Disruption of retromer components such as VPS-35, or early endosomal regulators including RME-8 and VPS-34, results in ZIP-3 degradation and robust induction of ATFS-1–dependent target genes, including those with extracellular functions. Importantly, this activation occurs independently of mitochondrial damage, revealing a signaling pathway in which endosomal dysfunction and stalled cargo recycling of cell surface receptors serve as primary triggers. We propose that endosomal swelling and impaired membrane flux are sensed upstream of ZIP-3, initiating a compensatory transcriptional response that preserves extracellular proteostasis and organismal health.

Mechanistically, we show that ZIP-3 stability is controlled by post-translational mechanisms involving ubiquitin-mediated degradation and kinase-dependent phosphorylation. Loss of surface receptors or downstream kinase activity markedly reduces ZIP-3 abundance, indicating that extracellular signaling actively maintains ZIP-3 as a repressor of UPR^EC^ under basal conditions. Under conditions of extracellular protein aggregation or trafficking failure, this repression is relieved through ZIP-3 turnover mediated by the E3 ubiquitin ligase WWP-1. Notably, phosphorylation of ZIP-3 confers protection from degradation, suggesting a regulatory switch by which extracellular cues calibrate UPR^EC^ activation.

The functional consequences of UPR^EC^ activation extend to organismal proteostasis and longevity. ATFS-1 activation, or genetic inhibition of ZIP-3, enhances extracellular proteome integrity and significantly improves healthspan in models expressing aggregation-prone proteins such as Aβ or TTR. Rescue of neuronal and muscular dysfunction in these models highlights UPR^EC^ as a protective pathway that buffers extracellular proteotoxic stress and mitigates tissue decline. These findings suggest that controlled activation of ATFS-1–dependent UPR^EC^ may represent a conserved adaptive strategy to counteract extracellular aggregation during aging.

Together, our findings determine an extracellular unfolded protein response (UPR^EC^) that links endosomal trafficking defects to transcriptional programs that preserve extracellular proteostasis. By identifying ZIP-3 degradation as a molecular switch that activates ATFS-1–dependent gene expression, this work reveals how cells sense disruptions in membrane trafficking and extracellular protein homeostasis to initiate adaptive protective responses. Because extracellular protein aggregation and endosomal dysfunction are conserved features of aging and neurodegenerative disorders such as Alzheimer’s Disease, the mechanisms uncovered here suggest that related surveillance pathways may operate in higher organisms to coordinate extracellular protein quality control. Elucidating whether analogous signaling networks involving mammalian ATF family transcription factors^46,47^ regulate extracellular proteostasis may therefore provide new insights into the pathogenesis of protein aggregation diseases and identify strategies to enhance extracellular proteome resilience during aging.

## Materials and Methods

### Worms, plasmids, and bacteria

N2 (wildtype), VC4372 W01A8.6 *(gk5453),* DCD23 LBP-2::tagRFP, OH906 otIs39 [unc-47(delta)::GFP + lin-15(+)], and BZ555 *dat-1_pr_::GFP* strains were obtained from the *Caenorhabditis* Genetics Center (Minneapolis, MN). LSD2104 *xchIs015* [pLSD134-P*hsp-16.2*::ssSel1:FLAG::superfolderGFP::spacer::humanAmyloidBeta1-42::let-858–3’UTR; pRF4 *rol-6(su1006)*] was obtained from the Ewald lab^27^. JKM2 *Is* [*rgef-1*p::Signalpeptide-Abeta(1-42)::*hsp-3*(IRES)::wrmScarlet-Abeta(1-42)::*unc-54*(3’UTR) + *rps-0*p::HygroR] and JKM7 *Is* [*myo-3*p::Signalpeptide-Abeta(1-42)::*hsp-3*(IRES)::wrmScarlet-Abeta(1-42)::*unc-54*(3’UTR) + *rps-0*p::HygroR] strains were obtained from the Kirstein lab^29^. SEE037 *scrIs008[unc-54p::hTTR(WT) + rol-6]*, SEE034 *uthIs378[unc-54p::hTTR(V30M) + rol-6]*, SEE106 *geIs101[rol-6(su1006)]*; *scrIs010[des-2p::myr::GFP + unc-122p::DsRed]* and SEE145 *uthIs378[unc-54p::hTTR(V30M) + rol-6]; scrIs010[des-2p::myr::gfp + unc-122p::DsRed*strains were obtained from the Encalada lab^28^. *fshr-1p::fshr-1::SL2::mKate* strain was obtained from the Beets lab^48^. mRuby2::RAB-11.1 strain was obtained from Douglas’ lab^49^.

ZIP-3-3x-FLAG and ZIP-3(RAAA) strains were generated via CRISPR-Cas9 in wildtype worms as described ^12^. The crRNAs (IDT) were co-injected with purified Cas9 protein, tracrRNA (Dharmacon), repair templates (IDT), and the *pRF4::rol-6(su1006)* plasmid as described ^50^. The *pRF4::rol-6 (su1006)* plasmid was a gift from Craig Mello ^50^. The *vha-6_pr_::daf-2::GFP* plasmid was gifted by Tian Xia and Anbing Shi labs^51^.

The *W01A8.6_pr_::GFP* plasmid was generated using conventional cloning of the *W01A8.6* promoter sequence and a portion of the first exon (2021 bp) into the pPD95.75 vector. The *F54E2.1_pr_::GFP* plasmid was generated using conventional cloning of the *F54E2.1* promoter sequence and the full first exon (4025 bp) into the pPD95.75 vector. The *myo-3_pr_::LBP-2-GFP* plasmid was generated using conventional cloning of the LBP-2 coding sequence (584 bp) into the pPD136.64 vector in frame with the *myo-3* promoter. Worms were raised with the HT115 strain of Escherichia coli, and RNAi was performed as described.

### Analysis of worm development

Worms were synchronized via bleaching and allowed to develop on HT115 bacteria plates at 20°C. The developmental stage is quantified as a percentage of the total number of animals that turned adult after an incubation of 58 hrs at 20°C as previously described^52^. Each experiment was performed three times.

### Lifespan assay

Lifespan analysis was carried out following an established protocol ^53,54^. Each strain was repeated at least twice. At least 50 animals were used per condition, and worms were scored for viability every second day, from day 1 of adulthood (treating the pre-fertile day preceding adulthood as t = 0). Young adult worms were transferred to fresh plates every other day, and the number of dead worms was recorded as events scored. Animals that were lost or burrowed in the medium, exhibiting protruding vulva (intestine protrudes from the vulva), or undergoing bagging (larvae hatching inside the worm body) were censored. Prism 8 (GraphPad) software was used for statistical analysis, and *p-values* were calculated using the log-rank (Kaplan-Meier) method.

### Protein analysis and antibodies

Synchronized worms were raised on plates with control(RNAi) or *cco-1*(RNAi) to the L4 stage before harvesting. The whole worm lysate preparation was previously described ^55^. Antibodies against β-actin (cell signaling), anti-FLAG M2 antibody (Sigma, F1804), RFP antibody (Rockland, 600-401-379), GFP antibody (Abcam, AB 183734), anti-phosphoserine/threonine antibody (BD Biosciences, 612548), and anti-phosphotyrosine antibody (Cell signaling, 8954S). All antibodies were diluted 1:2000. Immunoblots were imaged using the ChemiDoc XRS+ system (Bio-Rad). All western blot experiments were performed at least three times.

### Immunoprecipitation

Synchronized worms were raised on plates with wildtype (N2) or ZIP-3-3xFLAG strains to the L4 stage before harvesting. The whole worm lysate preparation was previously described ^55^ with a protease inhibitor and phosphatase inhibitor cocktail.

Anti-FLAG magnetic beads were gently resuspended by pipetting, and 10 µL of bead suspension was transferred to a microcentrifuge tube. The beads were washed twice with 0.5 mL TBS buffer (50 mM Tris-HCl, 150 mM NaCl, pH 7.4) by gentle pipetting and magnetic separation. Washed beads were incubated with 500 µL of cell lysate for 2 h at room temperature or overnight at 4 °C with gentle rotation to allow binding of ZIP-3-3xFLAG. Anti-FLAG M2 Magnetic beads (Sigma, M8823-1mL) are used for Immunoprecipitation using the ZIP-3-3xFLAG strain. Following incubation, beads were separated magnetically, and the supernatant was collected to assess unbound protein. The beads were then washed three times with PBST (136.89 mM NaCl, 2.67 mM KCl, 8.1 mM Na_₂_HPO_₄_, 1.76 mM KH_₂_PO_₄_, 0.5% Tween-20), with 5 min rotation for each wash. For denaturing elution, beads were resuspended in 50 µL of 1× protein sample loading buffer and boiled for 5 min. After magnetic separation, the eluates were analyzed by SDS-PAGE with anti-phosphoserine/threonine and anti-phosphotyrosine antibodies.

### TMRE staining

TMRE staining was performed by synchronizing worms and raising them on plates pre-soaked with S-Basal buffer containing TMRE at a final concentration of 100 μM (Sigma, 87917) ^56^. Before imaging, TMRE-stained worms were transferred to plates seeded with control (RNAi) bacteria and incubated for 3 h to eliminate TMRE-containing bacteria from the digestive tract. Images were acquired using a Zeiss LSM800 confocal microscope equipped with Airyscan, with identical exposure settings applied across all conditions. TMRE fluorescence analysis was performed as previously described. Briefly, average pixel intensity values were obtained by sampling images from individual worms, and the mean fluorescence intensity for each animal was quantified using ImageJ (http://rsb.info.nih.gov/ij/). Fluorescence intensity was measured from threshold-adjusted images for each condition using biological triplicates. Statistical comparisons were performed using Student’s *t*-test or one-way analysis of variance (ANOVA) followed by Tukey’s post hoc test, as appropriate.

### RNA isolation and qRT-PCR

RNA isolation and quantitative reverse transcriptase PCR (qRT-PCR) analysis were previously described ^57^. Worms were synchronized by bleaching, raised on HT115 *E. coli,* and harvested at the L4 stage. Total RNA was extracted from frozen worm pellets using Trizol reagent, and 500 ng RNA was used for cDNA synthesis with qScript™ cDNA SuperMix (QuantaBio). qPCR was performed using iQ™ SYBR® Green Supermix (Bio-Rad Laboratories). All qPCR results were repeated at least 3 times and performed in triplicate. A two-tailed Student’s t-test was employed to determine the level of statistical significance.

### Microscopy

*C. elegans* were imaged using either a Zeiss AxioCam 506 mono camera mounted on a Zeiss Axio Imager Z2 microscope or a Zeiss AxioCam MRc camera mounted on a Zeiss SteREO Discovery.V12 stereoscope. Images with high magnification (×63) were obtained using the Zeiss Apotome 2. Exposure times were the same in each experiment. Cell cultures were imaged with the Zeiss LSM800 microscope. With Zen 2.3 Blue software. All images are representative of more than three images. Quantification of fluorescent intensity as well as creating binary skeleton-like structures, was done with ImageJ.

### Gene set enrichment analysis

The gene set was downloaded from the WormBase Ontology Browser. mRNA abundance was measured and ranked by reads per kilobase per million reads from RNA-seq data. Pre-ranked gene set enrichment analysis was performed with GSEA3.0 software with ‘classical’ scoring.

### Statistics

All experiments were performed at least three times, yielding similar results, and comprised biological replicates. The sample size and statistical tests were chosen based on previous studies with similar methodologies, and the data met the assumptions for each test. No statistical method was used in deciding sample sizes. No blinded experiments were performed, and randomization was not used. For all figures, the mean ± SD is represented unless otherwise noted. Prism 8 (GraphPad) is used for statistical analysis and graph creation.

## Acknowledgements.

We thank Dr. Collin Ewald, Dr. Janine Kirtstein, Dr. Sandra Encalada, Dr. Peter Douglas, and Dr. Isabel Beets for sharing some worm strains with us. We thank Dr. Barth Grant for his advice on and for sharing the endosomal recycling reporter worm strains. We also thank Dr. Hira Goel for his advice and guidance throughout the project. We thank the *Caenorhabditis* Genetics Center for providing *C. elegans* strains, funded by NIH Office of Research 362 Infrastructure Programs (P40OD010440). This work was supported by the National Institutes of Health grants (R01AG040061, R01AG047182, and R37-AG047182-07) to C.M.H. and the Natural Sciences and Engineering Research Council (NSERC) postdoctoral fellowship to A.M. The authors are solely responsible for the content.

## Author’s contributions

A.M. and C.M.H. planned the experiments. A.M. and Y.D. generated worm strains. A.M. performed experiments. A.M. and C.M.H. wrote the manuscript.

## Conflict of Interest

The authors declare no conflicts of interest.

## Supplementary Figures

**Supplementary Figure 1.**
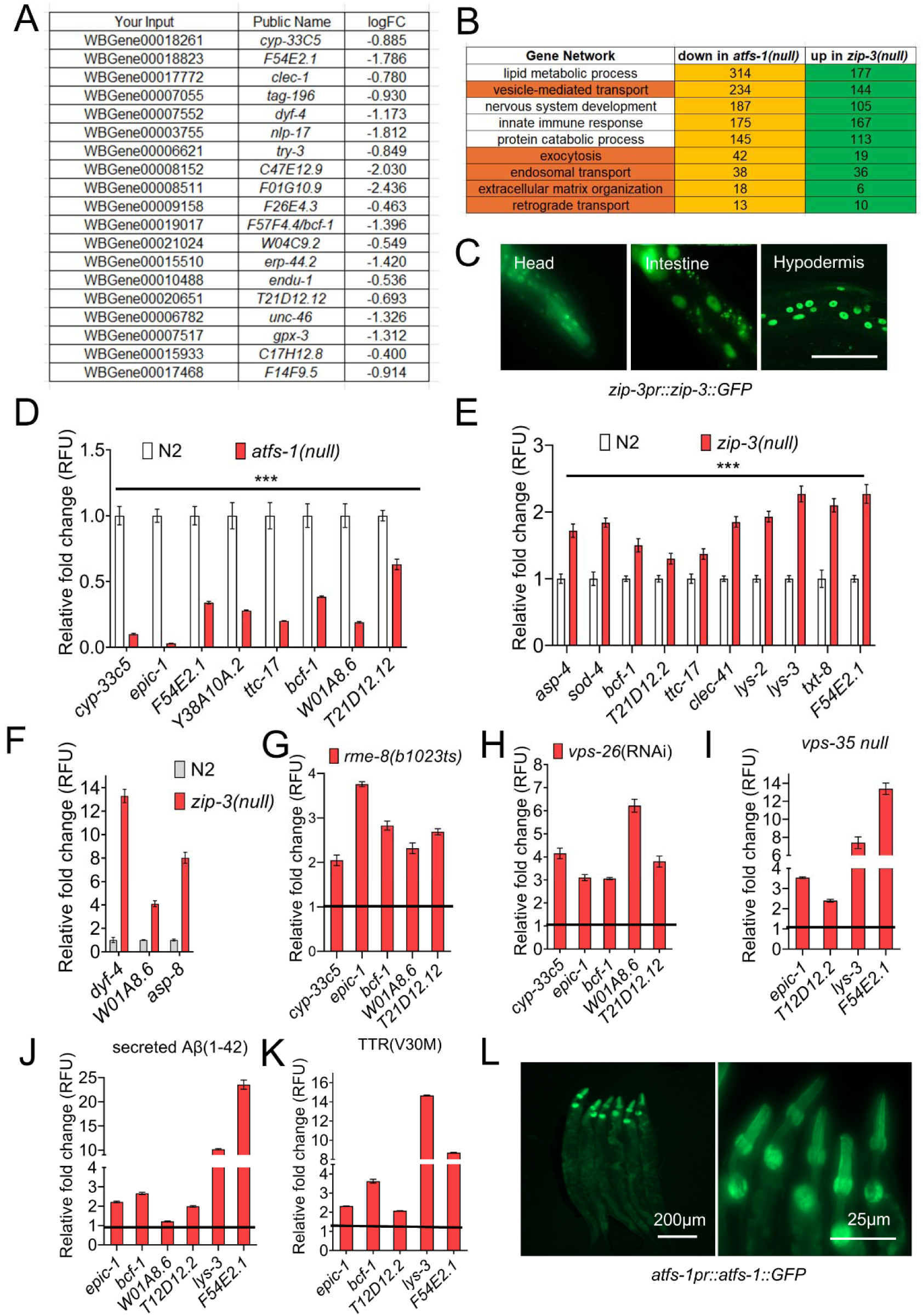
ATFS-1 promotes expression of endosomal recycling components and extracellular genes that limit protein aggregation. **A.** List of extracellular proteases and chaperones that regulate extracellular protein aggregation. **B.** Representative fluorescent image of *zip-3_pr_::zip-3-gfp* animals at the young adult stage. *zip-3* is expressed ubiquitously throughout the body, with notable high expression in the head, intestine, and hypodermis of animals. In the head (pharynx) *zip-3* is expressed in numerous neuronal cell bodies. Scale bar 50µm. **C.** Gene enrichment analysis of genes that are reduced in *atfs-1(null)* and increased in *zip-3(null); spg-7*(RNAi) worms. Dark orange color shows the categories vital to regulate extracellular proteostasis. **D.** Quantification of mRNA transcripts of extracellular genes (*cyp-33c5, epic-1, F54E2.1, Y38A10A.2, ttc-17, bcf-1, W01A8.6*, and *T21D12.12*) by qPCR in wildtype and *atfs-1(null)* at the L3 larval stage. N = 3, biologically independent samples. \*\*\**p* < 0.001 (one-way ANOVA). **E-F.** Quantification of mRNA transcripts of extracellular genes (*asp-4, sod-4*, *ttc-17, bcf-1, T21D12.12, clec-41, lys-2, lys-3, txt-8, F54E2.1, dyf-4, W01A8.6 and asp-8*) by qPCR in wildtype and *zip-3(null)* at L3 larval stage. N = 3, biologically independent samples. \*\*\**p* < 0.001 (one-way ANOVA). **G.** Bar graphs showing the expression of genes by qPCR in the *rme-1(b1023ts)* mutants compared to wildtype (RFU=1) at the L3 stage, grown at 15°C. **H.** Bar graphs showing the expression of genes by qPCR following control and *vps-26*(RNAi) (RFU=1) at the L3 stage. **I.** Bar graphs showing the expression of genes by qPCR in the *vps-35(ok1880)* mutants compared to wildtype (RFU=1) at the L3 stage. **J-K.** Bar graphs showing the expression of genes by qPCR in muscle secreted Aβ(1-42) and TTR(V30M) compared to wildtype (RFU=1) at the L3 stage. **G-K.** Graphs show the mean of two independent replicates with error bars showing the standard errors of the mean. \*\*\**p* < 0.001 (one-way ANOVA). **L.** Representative fluorescent image of *atfs-1_pr_::atfs-1-gfp* animals at the young adult stage (scale bar 200µm). *atfs-1* is expressed ubiquitously throughout the body, with notable high expression in the head of animals (scale bar 25µm).

**Supplementary Figure 2.**
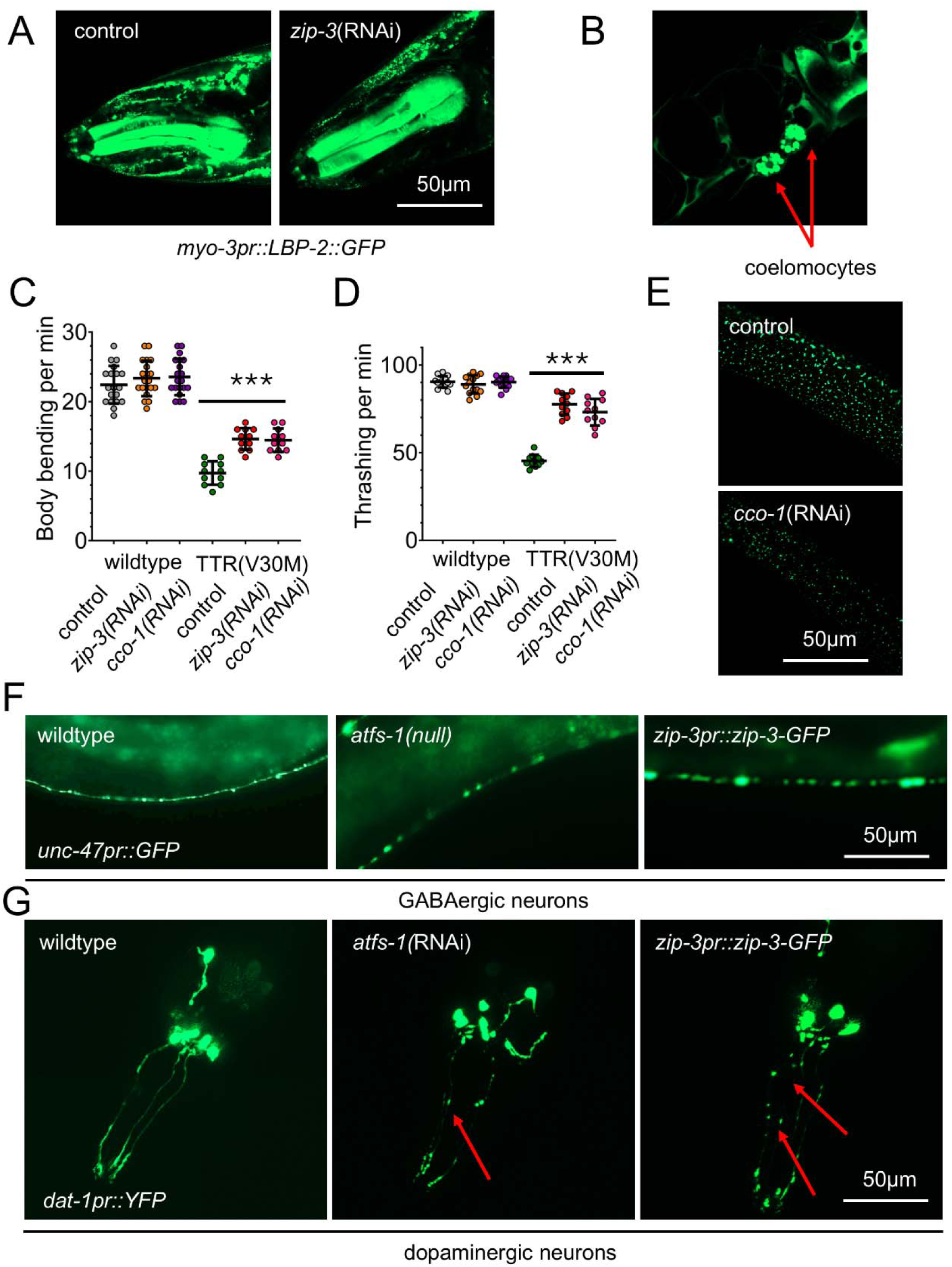
Inhibition of *atfs-1* and *zip-3* overexpression leads to neurodegeneration. **A.** Representative image of *myo-3_pr_::LBP-2-GFP* expression following control and *zip-3*(RNAi). Scale bar, 50μm. N =3, biologically independent replicates. **B.** Representative image of *myo-3_pr_::LBP-2-GFP* worms with GFP detected in the coelomocytes showing that the coelomocyte cells have endocytosed the extracellular LBP-2::GFP secreted from the muscle cells. **C-D.** Dot plots showing the rate of thrashing and body bending of wildtype and TTR(V30M) worms grown on control, *zip-3*(RNAi), and *cco-1*(RNAi) plates. Data represents the mean of three independent biological replicates with error bars showing the standard error of the mean. N = 3, biologically independent samples. \*\*\**p* < 0.001 (one-way ANOVA). **E.** Representative images of worms expressing *hsp-16.2_pr_::ssA*β*(1-42)-GFP* labelled protein aggregation puncta grown at 37°C for 1 hr during Day 1 adult following control, and *cco-1*(RNAi). Scale bar, 50μm. N =3, biologically independent replicates. **F.** Representative image of *unc-47_pr_::GFP* expression in the GABAergic neurons of wildtype, *atfs-1(null),* and *zip-3_pr_::zip-3-GFP* worms. Scale bar, 50μm. N =3, biologically independent replicates. **G.** Representative image of *dat-1_pr_::GFP* expression in the dopaminergic neurons of wildtype, *atfs-1(RNAi),* and *zip-3_pr_::zip-3-GFP* worms. Scale bar, 50μm. N =3, biologically independent replicates.

**Supplementary Figure 3.**
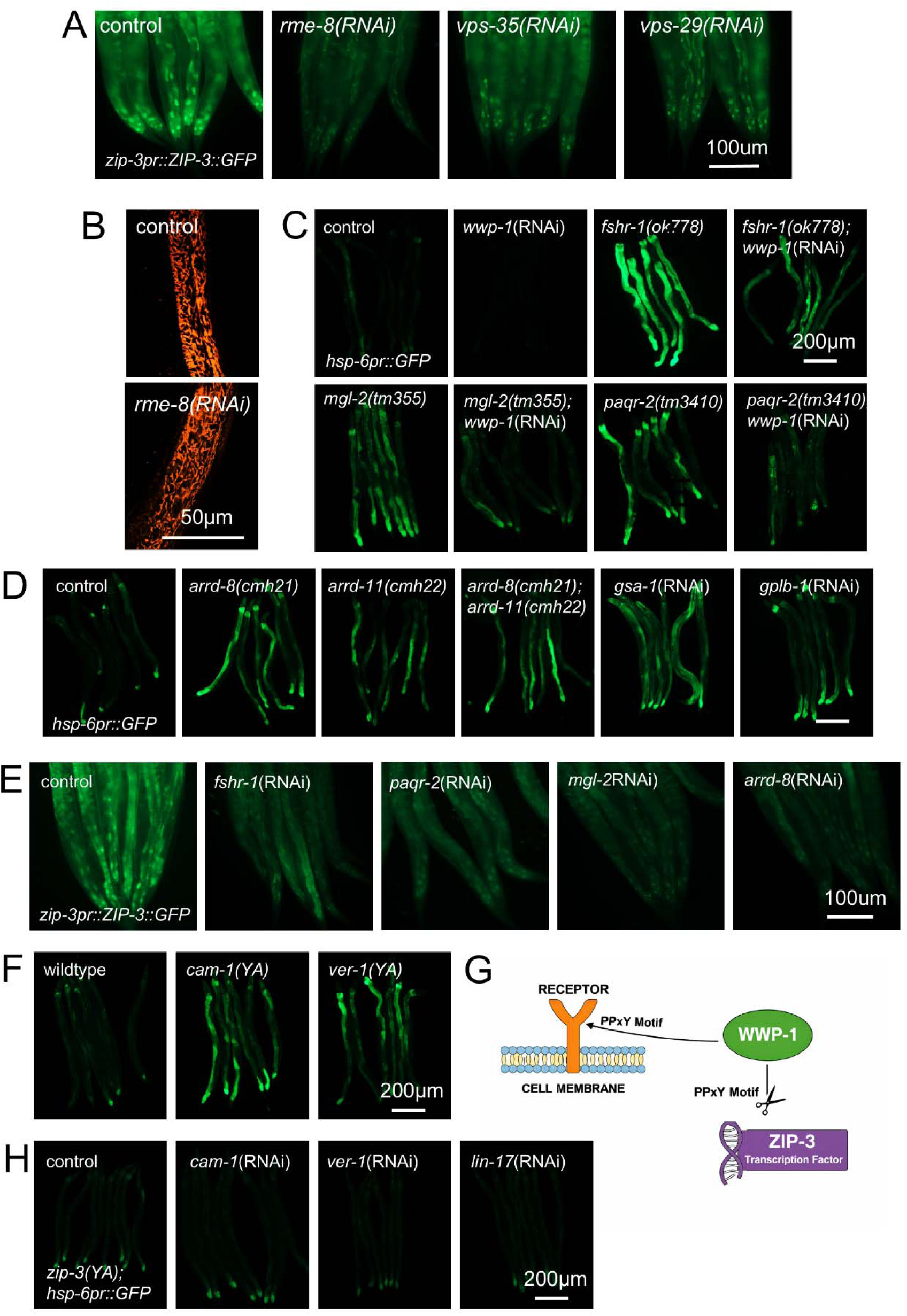
Inhibiting cell surface receptors leads to ZIP-3 degradation and ATFS-1 activation. **A.** Representative image of *zip-3_pr_::ZIP-3::GFP* expression following control, *rme-8*(RNAi), *vps-35*(RNAi), and *vps-29*(RNAi). Scale bar, 100μm. N =3, biologically independent replicates. **B.** Representative images of TMRE staining on wildtype worms at the L4 stage grown on control and rme-8(RNAi) HT115 bacteria. Scale bar, 50μm. N =3, biologically independent replicates. **C.** Representative image of UPR^mt^ reporter *hsp-6_pr_::GFP* expression in wildtype, *fshr-1(ok778), mgl-2(tm355),* and *paqr-2(tm3410)* worms grown on control and *wwp-1*(RNAi) HT115 bacteria. Scale bar, 200μm. N =3, biologically independent replicates. **D.** Representative image of UPR^mt^ reporter *hsp-6_pr_::GFP* expression in wildtype, *arrd-8(cmh21), arrd-11(cmh22), gsa-1*(RNAi), and *gplb-1*(RNAi) worms grown on control and *wwp-1*(RNAi) HT115 bacteria. Scale bar, 200μm. N =3, biologically independent replicates. **E.** Representative image of *zip-3_pr_::ZIP-3::GFP* expression following control, *fshr-1*(RNAi), *paqr-2*(RNAi), *mgl-2*(RNAi) and *arrd-8*(RNAi). Scale bar, 100μm. N =3, biologically independent replicates. **F.** Representative image of UPR^mt^ reporter *hsp-6_pr_::GFP* expression in wildtype, *cam-1(YA)* and *ver-1(YA)* worms. PPxY motifs on the CAM-1 and VER-1 genes have been mutated. Scale bar, 200μm. N =3, biologically independent replicates. **G.** A schematic showing WWP-1 interaction with its substrates, cell surface receptors, and ZIP-3. Lack of one substrate makes the interaction with the other substrate more prominent. **H.** Representative image of UPR^mt^ reporter *hsp-6_pr_::GFP* expression in ZIP-3(YA) mutants following control, *cam-1*(RNAi), *ver-1*(RNAi) and *lin-17*(RNAi). The PPxY motif of the ZIP-3 gene has been mutated. Scale bar, 200μm. N =3, biologically independent replicates.

**Supplementary Figure 4.**
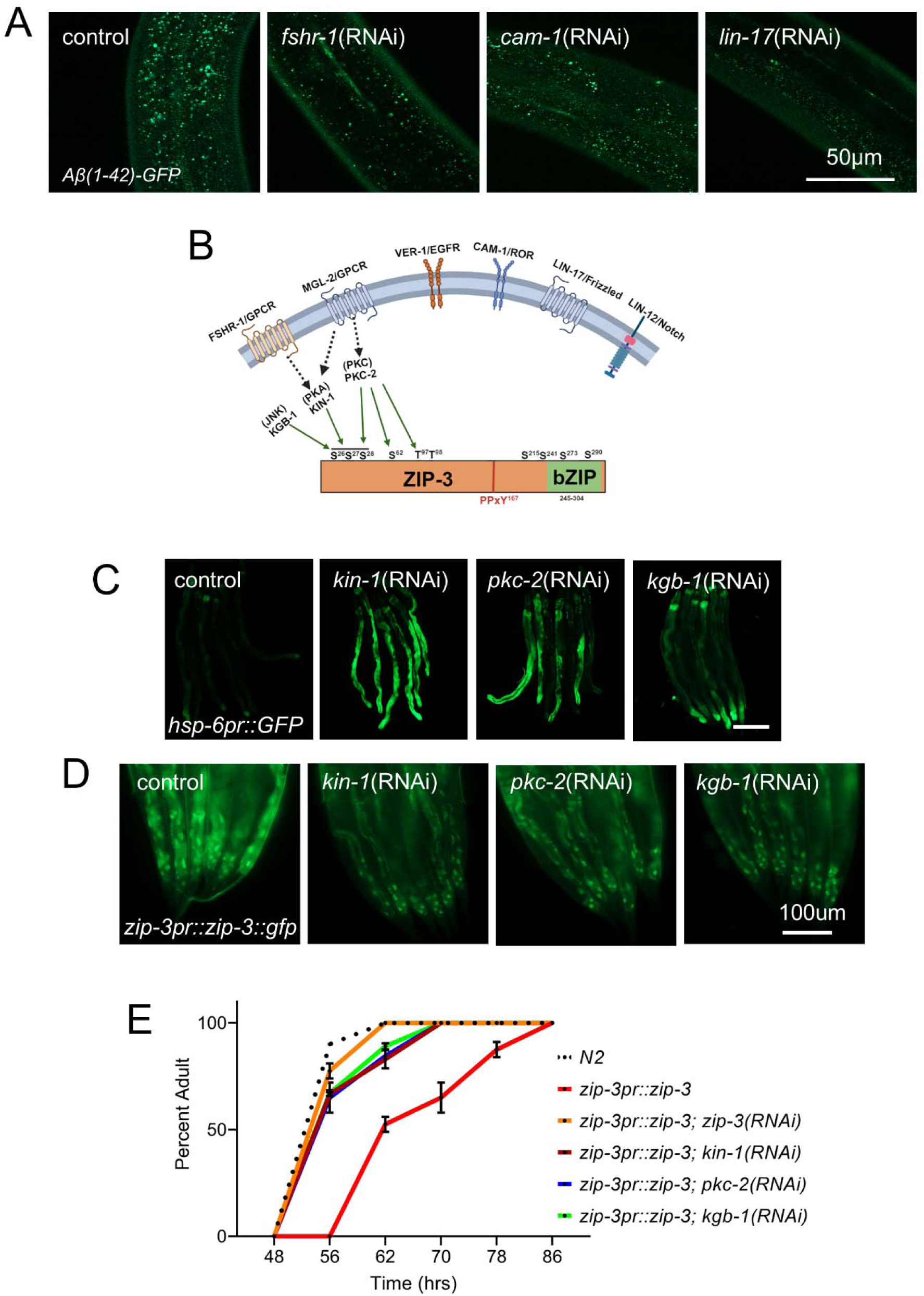
Inhibiting cell surface receptors limits Aβ(1-42) aggregation. **A.** Representative images of worms expressing *hsp-16.2_pr_::ssA*β*(1-42)-GFP* labelled protein aggregation puncta grown at 37°C for 1 hr during Day 1 adult following control, *fshr-1*(RNAi), *cam-1*(RNAi), and *lin-17*(RNAi). Scale bar, 50μm. N =3, biologically independent replicates. **B.** A schematic of the ZIP-3 protein showing the predicted phosphorylation sites by different families of kinases (JNK, PKA, and PKC). Ligand binding at the cell surface receptors leads to activation of downstream kinases and ZIP-3 phosphorylation, which in turn leads to ZIP-3 stability. **C.** Representative image of UPR^mt^ reporter *hsp-6_pr_::GFP* expression following control, *kin-1*(RNAi), *pkc-2*(RNAi), and *kgb-1*(RNAi). Scale bar, 200μm. N =3, biologically independent replicates. **D.** Representative image of *zip-3_pr_::ZIP-3::GFP* expression following control, *kin-1*(RNAi), *pkc-2*(RNAi), and *kgb-1*(RNAi). Scale bar, 100μm. N =3, biologically independent replicates. **E.** Developmental rate plotted as the time taken to become an adult from egg. The graph represents the mean of two independent biological replicates, and the error bars shows standard error of means. \*\*\**p* < 0.001 (one-way ANOVA), all RNAi conditions vs *zip-3_pr_::ZIP-3*.

**Supplementary Figure 5.**
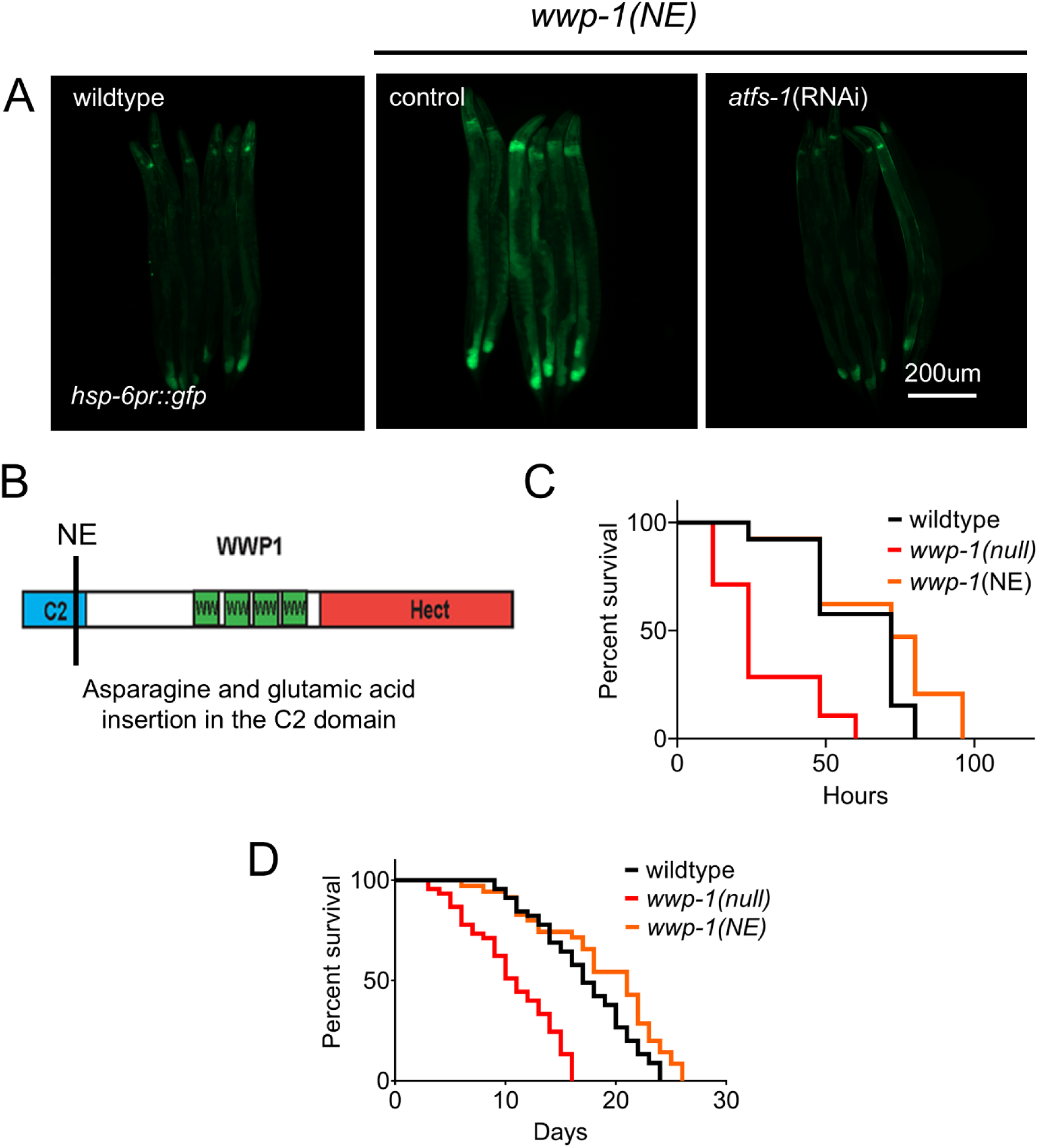
Mutating the C2 domain on WWP-1 activates ATFS-1-dependent transcription. **A.** Representative image of UPR^mt^ reporter *hsp-6_pr_::GFP* expression in wildtype and *wwp-1*(*NE)* mutants grown on control and *atfs-1*(RNAi). Scale bar, 200μm. N =3, biologically independent replicates. **B.** A schematic of the mutation on the C2 domain of WWP-1. The C2 domain is required to facilitate cell membrane tethering of WWP-1 and interaction with its HECT domain to promote autoinhibition. **C.** Survival graph of wildtype (N2), wwp-1(null) and wwp-1(NE) worms *on Pseudomonus aeruginosa* (PA14) at 25°C. **D.** Lifespan graphs for wildtype, *wwp-1*(null), and *wwp-1(NE)*. N =3, biologically independent replicates. \**p* < 0.01 (**C, D**) (log-rank test).

**Supplementary Figure 6.**
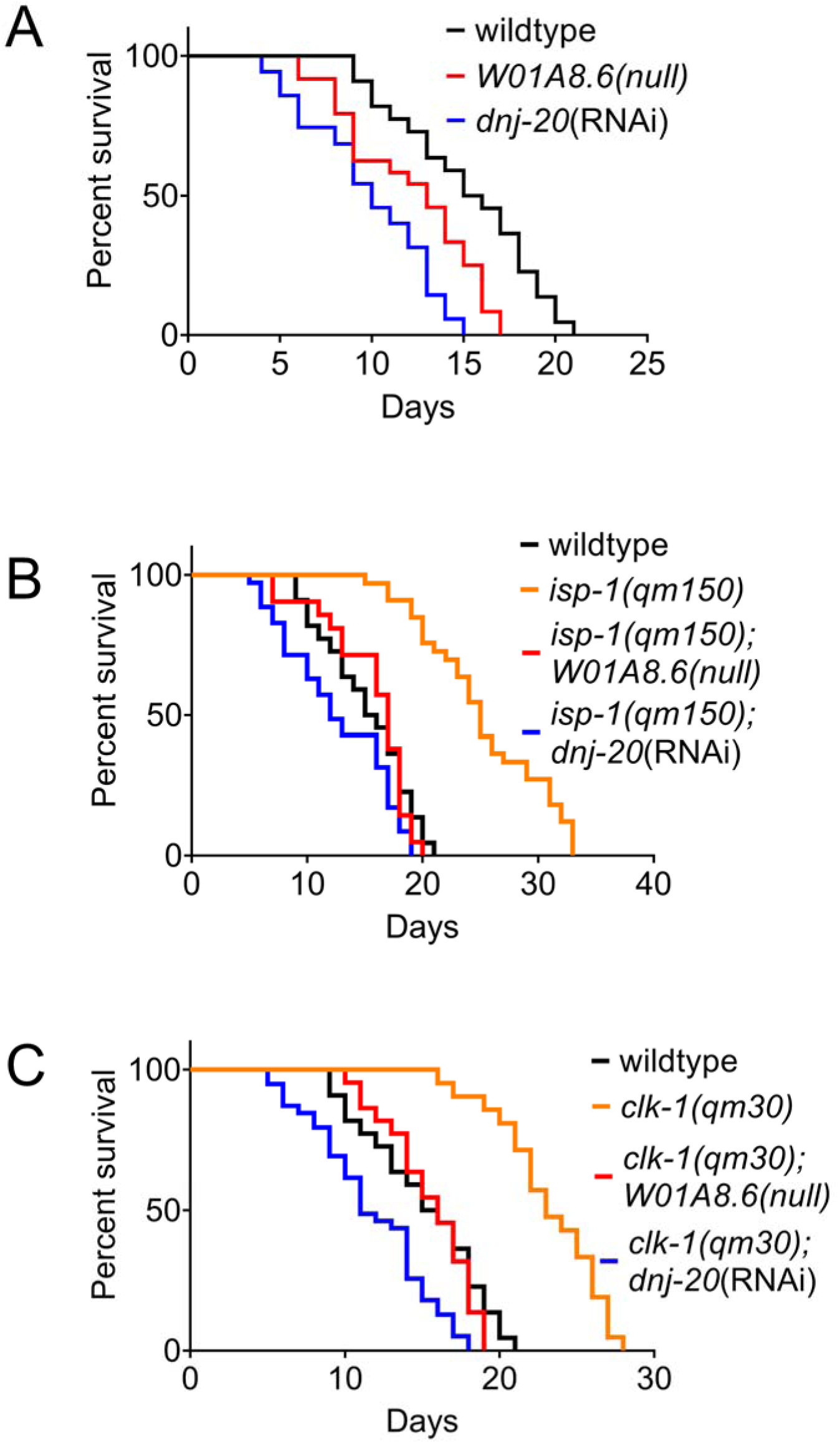
Inhibiting ATFS-1-dependent extracellular protease *W01A8.6* and extracellular chaperone DNJ-20 limits the long lifespan of *clk-1* and *isp-1* mutants. **A.** Lifespan graphs for wildtype, *W01A8.6(null),* and *dnj-20*(RNAi). N =2, biologically independent replicates. \**p* < 0.01 (wildtype vs *W01A8.6* and wildtype vs *dnj-20*) (log-rank test). **B.** Lifespan graphs for *isp-1(qm150)* mutants compared to *isp-1(qm150)*, *W01A8.6(null),* and *isp-1(qm150), dnj-20*(RNAi). N =2, biologically independent replicates. \*\**p* < 0.001 (*isp-1* vs *isp-1, W01A8.6* and *isp-1* vs *isp-1, dnj-20*) (log-rank test). **C.** Lifespan graphs for *clk-1(qm30)* mutants compared to *clk-1(qm30)*, *W01A8.6(null),* and *clk-1(qm30), dnj-20*(RNAi). N =2, biologically independent replicates. \*\**p* < 0.001 (*clk-1* vs *clk-1, W01A8.6* and *clk-1* vs *clk-1, dnj-20*) (log-rank test).

## Notes

### Competing Interest Statement

The authors have declared no competing interest.

## References

1. Walter, P. & Ron, D. The unfolded protein response: From stress pathway to homeostatic regulation. Science (80-.). 334, 1081–1086 (2011).

2. Higuchi-Sanabria, R., Frankino, P. A., Paul, J. W., Tronnes, S. U. & Dillin, A. A Futile Battle? Protein Quality Control and the Stress of Aging. Dev. Cell 44, 139–163 (2018).

3. Shpilka, T. & Haynes, C. M. The mitochondrial UPR: Mechanisms, physiological functions and implications in ageing. Nat. Rev. Mol. Cell Biol. 19, 109–120 (2018).

4. Vaquer-Alicea, J. & Diamond, M. I. Propagation of protein aggregation in neurodegenerative diseases. Annu. Rev. Biochem. 88, 785–810 (2019).

5. Gao, H. et al. Mitochondrial complex I deficiency induces Alzheimer’s disease–like signatures that are reversible by targeted therapy. Alzheimer’s Dement. 21, 1–19 (2025).

6. Stojakovic, A. et al. Partial inhibition of mitochondrial complex I ameliorates Alzheimer’s disease pathology and cognition in APP/PS1 female mice. Commun. Biol. 4, (2021).

7. Mathys, H. et al. Single-cell multiregion dissection of Alzheimer’s disease. Nature (2024). doi:10.1038/s41586-024-07606-7

8. Sorrentino, V. et al. Enhancing mitochondrial proteostasis reduces amyloid-β proteotoxicity. Nature 552, 187–193 (2017).

9. Nargund, A., Pellegrino, M. W., Fiorese, C. J., Baker, B. M. & Haynes, C. M. Mitochondrial Import Efficiency of ATFS-1 Regulates Mitochondrial UPR Activation. Science (80-.). 337, 587–590 (2012).

10. Nargund, A. M., Fiorese, C. J., Pellegrino, M. W., Deng, P. & Haynes, C. M. Mitochondrial and nuclear accumulation of the transcription factor ATFS-1 promotes OXPHOS recovery during the UPRmt. Mol. Cell 58, 123–133 (2015).

11. Shpilka, T. et al. UPRmt scales mitochondrial network expansion with protein synthesis via mitochondrial import in Caenorhabditis elegans. Nat. Commun. 12, (2021).

12. Deng, P. et al. Mitochondrial UPR repression during Pseudomonas aeruginosa infection requires the bZIP protein ZIP-3. Proc. Natl. Acad. Sci. U. S. A. 116, 6146–6151 (2019).

13. Reinke, A. W., Baek, J., Ashenberg, O. & Keating, A. E. Networks of bZIP Protein-Protein Interactions Diversified Over a Billion Years of Evolution. Science (80-.). 340, 730–735 (2013).

14. Szabo, M. P., Mishra, S., Knupp, A. & Young, J. E. The role of Alzheimer’s disease risk genes in endolysosomal pathways. Neurobiol. Dis. 162, 105576 (2022).

15. Young, J. E., Holstege, H., Andersen, O. M., Petsko, G. A. & Small, S. A. On the causal role of retromer-dependent endosomal recycling in Alzheimer’s disease. Nat. Cell Biol. 25, 1394–1397 (2023).

16. Cullen, P. J. & Steinberg, F. To degrade or not to degrade: mechanisms and significance of endocytic recycling. Nat. Rev. Mol. Cell Biol. 19, 679–696 (2018).

17. Shi, A. et al. Regulation of endosomal clathrin and retromer-mediated endosome to Golgi retrograde transport by the J-domain protein RME-8. EMBO J. 28, 3290–3302 (2009).

18. Sorkin, A. & Von Zastrow, M. Signal transduction and endocytosis: Close encounters of many kinds. Nat. Rev. Mol. Cell Biol. 3, 600–614 (2002).

19. Fletcher, K. A., Alkurashi, M. H. & Lindsay, A. J. Endosomal recycling inhibitors downregulate the androgen receptor and synergise with enzalutamide. Invest. New Drugs 42, 14–23 (2024).

20. Mishra, S. et al. The Alzheimer’s gene SORL1 is a regulator of endosomal traffic and recycling in human neurons. Cell. Mol. Life Sci. 79, 1–22 (2022).

21. Small, S. A. et al. Model-Guided Microarray Implicates the Retromer Complex in Alzheimer ‘ s Disease. 909–919 (2005). doi:10.1002/ana.20667

22. Kachergus, J. M. et al. VPS35 Mutations in Parkinson Disease. 162–167 (2011). doi:10.1016/j.ajhg.2011.06.001

23. Tang, F., Liu, W., Hu, J., Mei, L. & Xiong, W. VPS35 Deficiency or Mutation Causes Dopaminergic Neuronal Loss by Impairing Mitochondrial Fusion and Article VPS35 Deficiency or Mutation Causes Dopaminergic Neuronal Loss by Impairing Mitochondrial Fusion and Function. CellReports 12, 1631–1643 (2015).

24. Gallotta, I. et al. Extracellular proteostasis prevents aggregation during pathogenic attack. Nature 584, 410–414 (2020).

25. Braun, J. E. A. Extracellular chaperone networks and the export of J-domain proteins. J. Biol. Chem. 299, 102840 (2023).

26. Durieux, J., Wolff, S. & Dillin, A. The cell-non-autonomous nature of electron transport chain-mediated longevity. Cell 144, 79–91 (2011).

27. Jongsma, E., Goyala, A., Mateos, J. M. & Ewald, C. Y. Removal of extracellular human amyloid beta aggregates by extracellular proteases in C. elegans. Elife 12, 1–21 (2023).

28. Madhivanan, K. et al. Cellular clearance of circulating transthyretin decreases cell-nonautonomous proteotoxicity in Caenorhabditis elegans. Proc. Natl. Acad. Sci. U. S. A. 115, E7710–E7719 (2018).

29. Gallrein, C. et al. Novel amyloid-beta pathology C. elegans model reveals distinct neurons as seeds of pathogenicity. Prog. Neurobiol. 198, 101907 (2021).

30. Wang, C. et al. A neurotransmitter atlas of C . elegans males and hermaphrodites. 1–46 (2024).

31. Robinson, S. B. et al. Molecular and Cellular Neuroscience Sequence determinants of the Caenhorhabditis elegans dopamine transporter dictating in vivo axonal export and synaptic localization. Mol. Cell. Neurosci. 78, 41–51 (2017).

32. Sato, K., Norris, A., Sato, M. & Grant, B. D. C . elegans as a model for membrane traffic *. (2014). doi:10.1895/wormbook.1.77.2

33. Rostislavleva, K. et al. Structure and flexibility of the endosomal Vps34 complex reveals the basis of its function on membranes. 7365, (2015).

34. Wallroth, A. & Haucke, V. Phosphoinositide conversion in endocytosis and the endolysosomal system. 293, 1526–1535 (2018).

35. Zhi, X. & Chen, C. WWP1: A versatile ubiquitin E3 ligase in signaling and diseases. Cell. Mol. Life Sci. 69, 1425–1434 (2012).

36. Sardana, R. & Emr, S. D. Membrane Protein Quality Control Mechanisms in the Endo-Lysosome System. Trends Cell Biol. 31, 269–283 (2021).

37. Sun, T., Wang, X., Lu, Q., Ren, H. & Zhang, H. CUP-5, the C. elegans ortholog of the mammalian lysosomal channel protein MLN1/TRPML1, is required for proteolytic degradation in autolysosomes. Autophagy 7, 1308–1315 (2011).

38. Blom, N., Sicheritz-pontén, T., Gupta, R., Gammeltoft, S. & Brunak, S. Prediction of post-translational glycosylation and phosphorylation of proteins from the amino acid sequence. Proteomics 4, 1633–1649 (2004).

39. Wiesner, S. et al. Autoinhibition of the HECT-Type Ubiquitin Ligase Smurf2 through Its C2 Domain. Cell 130, 651–662 (2007).

40. Courivaud, T. et al. Functional Characterization of a WWP1 / Tiul1 Tumor-derived Mutant Reveals a Paradigm of Its Constitutive Activation in Human Cancer *. 290, 21007–21018 (2015).

41. Feng, J., Bussière, F. & Hekimi, S. Mitochondrial Electron Transport Is a Key Determinant of Life Span in Caenorhabditis elegans. Dev. Cell 1, 633–644 (2001).

42. Felkai, S. et al. CLK-1 controls respiration, behavior and aging in the nematode Caenorhabditis elegans. EMBO J. 18, 1783–1792 (1999).

43. Dillin, A. et al. Rates of behavior and aging specified by mitochondrial function during development. Science (80-.). 298, 2398–2401 (2002).

44. Sutherland, T. E., Dyer, D. P. & Allen, J. E. The extracellular matrix and the immune system: A mutually dependent relationship. Science (80-.). 379, (2023).

45. Bandzerewicz, A. & Gadomska-Gajadhur, A. Into the Tissues: Extracellular Matrix and Its Artificial Substitutes: Cell Signalling Mechanisms. Cells 11, (2022).

46. Fiorese, C. J. et al. The Transcription Factor ATF5 Mediates a Mammalian Mitochondrial UPR. Curr. Biol. 26, 2037–2043 (2016).

47. Anderson, N. S. & Haynes, C. M. Folding the Mitochondrial UPR into the Integrated Stress Response. Trends Cell Biol. 30, 428–439 (2020).

48. Kenis, S. et al. Ancestral glycoprotein hormone-receptor pathway controls growth in C . elegans. 1–15 (2023). doi:10.3389/fendo.2023.1200407

49. Watterson, A. et al. Intracellular lipid surveillance by small G protein geranylgeranylation. Nature 605, 736–740 (2022).

50. Dokshin, G. A., Ghanta, K. S., Piscopo, K. M. & Mello, C. C. Robust genome editing with short single-stranded and long, partially single-stranded DNA donors in caenorhabditis elegans. Genetics 210, 781–787 (2018).

51. Chen, D. et al. Revealing Functional Crosstalk between Distinct Bioprocesses through Reciprocal Functional Tests of Genetically Interacting Genes Article Revealing Functional Crosstalk between Distinct Bioprocesses through Reciprocal Functional Tests of Genetically Interacting Genes. 2646–2658 (2019). doi:10.1016/j.celrep.2019.10.076

52. Mallick, A. et al. AMP accumulation during mitochondrial stress induces transcription of cytosolic and mitochondrial protein synthesis components via NHR-180. 1, (2025).

53. Amrit, F. R. G., Ratnappan, R., Keith, S. A. & Ghazi, A. The C. elegans lifespan assay toolkit. Methods 68, 465–475 (2014).

54. Mallick, A., Ranawade, A., van den Berg, W. & Gupta, B. P. Axin-Mediated Regulation of Lifespan and Muscle Health in C. elegans Requires AMPK-FOXO Signaling. iScience 23, 101843 (2020).

55. Yoneda, T. et al. Compartment-specific perturbation of protein handling activates genes encoding mitochondrial chaperones. J. Cell Sci. 117, 4055–4066 (2004).

56. Mallick, A. & Haynes, C. M. Methods to analyze the mitochondrial unfolded protein response (UPRmt). Methods Enzymol. 707, 543–564 (2024).

57. Lin, Y. F. et al. Maintenance and propagation of a deleterious mitochondrial genome by the mitochondrial unfolded protein response. Nature 533, 416–419 (2016).

